# Expression and manufacturing of protein therapeutics in spirulina

**DOI:** 10.1101/2021.01.25.427910

**Authors:** Benjamin Jester, Hui Zhao, Mesfin Gewe, Thomas Adame, Lisa Perruzza, David Bolick, Jan Agosti, Nhi Khuong, Rolf Kuestner, Caitlin Gamble, Kendra Cruickshank, Jeremy Ferrara, Rachelle Lim, Troy Paddock, Colin Brady, Stacey Ertel, Mia Zhang, Michael Tasch, Tracy Saveria, David Doughty, Jacob Marshall, Damian Carrieri, Jamie Lee, Lauren Goetsch, Jason Dang, Nathaniel Sanjaya, David Fletecher, Anissa Martinez, Bryce Kadis, Kristjian Sigmar, Esha Afreen, Tammy Nguyen, Amanda Randolph, Alexandria Taber, Ashley Krzeszowski, Brittney Robinett, Fabio Grassi, Richard Guerrant, Michael Spigarelli, Ryo Takeuchi, Brian Finrow, Craig Behnke, James Roberts

## Abstract

*Arthrospira platensis* (commonly known as spirulina) is a photosynthetic cyanobacterium^1^. It is a highly nutritious food that has been consumed for decades in the US, and even longer by indigenous cultures^2^. Its widespread use as a safe food source and proven scalability have driven frequent attempts to convert it into a biomanufacturing platform. But these were repeatedly frustrated by spirulina’s genetic intractability. We report here efficient and versatile genetic engineering methodology for spirulina that allows stable expression of bioactive protein therapeutics at high levels. We further describe large-scale, indoor cultivation and downstream processing methods appropriate for the manufacturing of biopharmaceuticals in spirulina. The potential of the platform is illustrated by pre-clinical development and human testing of an orally delivered antibody therapeutic against campylobacter, a major cause of infant mortality in the developing world and a growing antibiotic resistance threat^3,4^. This integrated development and manufacturing platform blends the safety of food-based biotechnology with the ease of genetic manipulation, rapid growth rates and high productivity characteristic of microbial platforms. These features combine for exceptionally low-cost production of biopharmaceuticals to address medical needs that are unfeasible with current biotechnology platforms.

## INTRODUCTION

The biotechnology industry has its origins in the domestication of cells as biological factories. Over the past 50 years this industry has been transformed by the discovery of methods for genetic engineering, which led to adoption of a succession of platform organisms for manufacturing recombinant biological products^5–7^. *E. coli* was the first, which was used to manufacture relatively small and simple therapeutic proteins^8,9^. Other complementary platform organisms were introduced in the 1980s and 1990s, most notably yeasts and mammalian cells. These did not replace earlier platforms, but rather enabled biomanufacturing to address medical problems those prior platforms had failed to solve. The advent of yeast genetic engineering made available a new suite of important protein products with more complex structures and post-translational modifications^10^. Mammalian cell transformation technologies soon followed, allowing the manufacturing of yet more complex proteins—for example, ones whose functions require mammalian glycosylation patterns—and also more efficient manufacturing of secreted proteins^11,12^.

Widespread adoption of new cellular platforms has been driven largely by 3 factors: the discovery of effective methods for their genetic manipulation; biological traits that facilitate the manufacture of products that are more difficult or impossible to produce with pre-existing platforms; and the development of robust, large-scale manufacturing technologies. Genetically engineered plants—particularly food crops—have long promised to provide a new platform for manufacturing biopharmaceutical products, with performance characteristics orthogonal to the traditional platforms mentioned above^13–19^. But this promise has not been realized for many reasons, including cumbersome genetic methods, slow growth rates, low product yields and regulatory constraints^20–22^.

Though not a true plant, photosynthetic spirulina is the only microorganism that is farmed as a food at commercial scale, worldwide. It has many unique attributes that differentiate it from existing expression platforms, which include simple, low-cost growth and downstream processing; photosynthetic metabolism; and the built-in safety afforded by manufacturing and delivering edible protein therapeutics in a food. In addition, its high protein content (50-70% of biomass) exceeds all other staple food crops, making it a strong candidate for the expression of therapeutic proteins. Spirulina’s asexual reproduction mitigates the risk of gene escape into the food chain, and the associated food security concerns and regulatory burden^23^. Spirulina therefore promises all the benefits of plant-based biopharmaceuticals, and—with the discoveries reported here—overcomes the challenges and limitations associated with commercial adoption of other food-based platforms.

We describe below our discovery of versatile genetic engineering methods for spirulina and development of indoor cultivation technology suitable for the large-scale manufacturing of biopharmaceuticals. We also report the development of an edible antibody-based therapeutic targeting gastrointestinal infection by *Campylobacter jejuni* to illustrate application of the spirulina platform to an important unmet medical need, including validation in animal models of campylobacteriosis, large-scale production under current Good Manufacturing Practices (cGMP), and evaluation of its safety and pharmacokinetics in a Phase 1 clinical trial.

## METHODS

See Supplemental Materials.

## RESULTS

### Spirulina are naturally competent for transformation

Our experimental analyses and methodological applications showed that spirulina are naturally competent for transformation despite being widely viewed as refractory to genetic manipulation^24–26^. Efficient transformation was achieved by co-cultivation of spirulina with companion microorganisms that induced competence (see below). Competent spirulina (UTEX LB1926 and NIES-39) were exposed to an integrating circular DNA vector containing an antibiotic-resistance marker and a gene of interest flanked on both sides by sequences homologous to the spirulina chromosome. Spirulina were then maintained in liquid culture under antibiotic selection. Microscopic clusters of green cells were detected after 2 weeks of cultivation, and after 3 weeks fully green cultures were apparent. Precise integration of the transforming DNA into the spirulina genome by homologous recombination was demonstrated by PCR amplification and DNA sequencing of chromosomal DNA using primers flanking the insertion site (Figures 1A, 1B). This transformation method yielded a pool of approximately 100 independent transformants, as determined by next-generation sequencing of a culture that had been exposed to a library of bar-coded, but otherwise identical, integrating DNA vectors. Consistent with these observations, the 15 genes associated with the pathway for natural competence in cyanobacteria^27^ are present as complete open reading frames in all available arthrospira genomes, including UTEX LB1926 and NIES-39 (Supplemental Figure S1).

**Figure 1:**
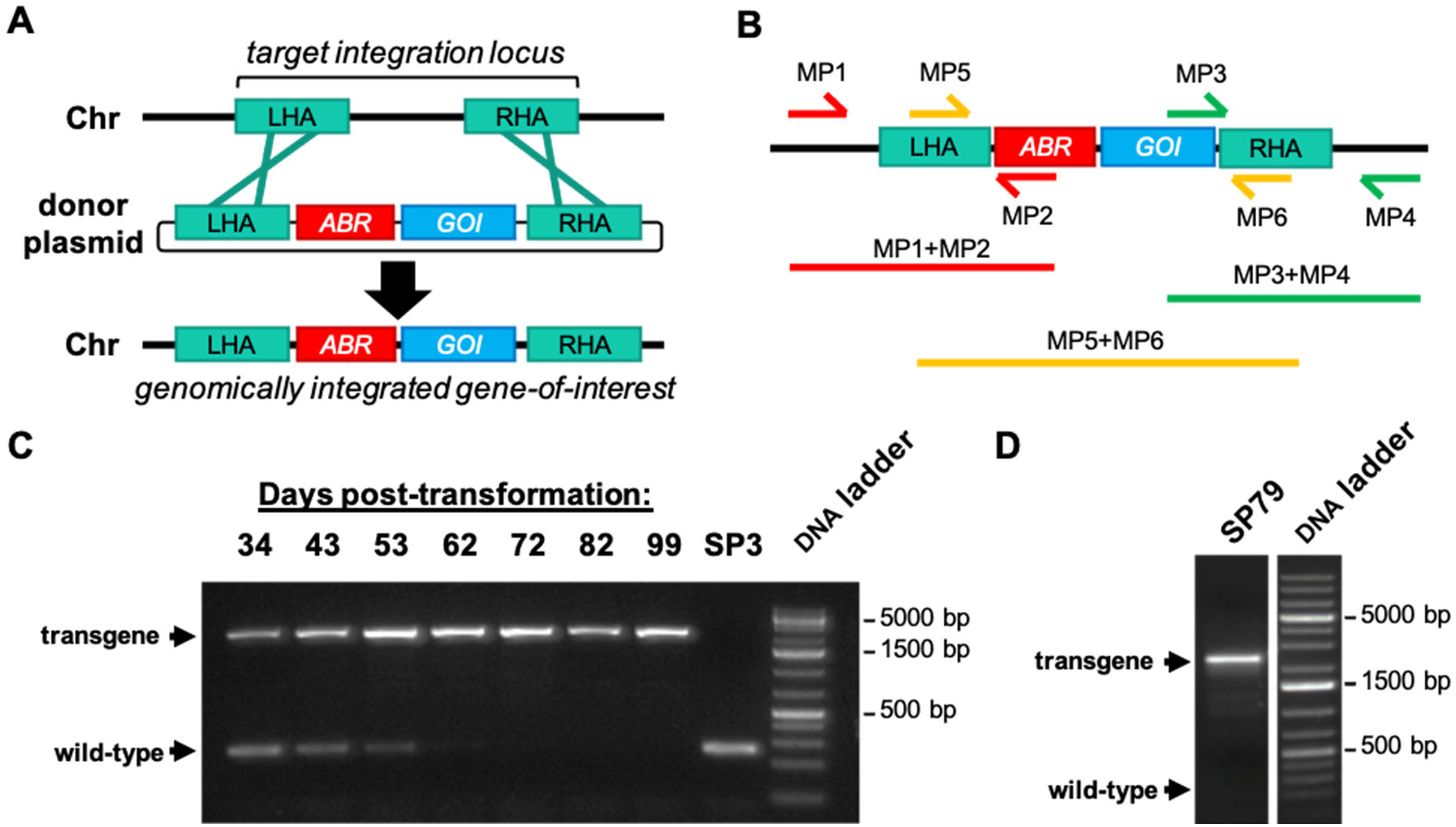
Homologous recombination of DNA into the spirulina chromosome (Chr). **A.** Illustration of genomic integration. Donor plasmid DNA containing an antibiotic-resistance (ABR) gene and a recombinant gene-of-interest (GOI) flanked by left and right homology arms (LHA and RHA respectively) is transformed into spirulina. A double-crossover event allows the ABR and GOI to be inserted at the target integration locus, replacing the intervening genomic DNA. **B.** Diagram of primer pairs used in PCR to genotype genomic integration. The primer pairs for amplification of the LHA and RHA include 1 priming site (MP1 and MP4) that is only present in the spirulina genome at the target locus and absent from the donor plasmid. The PCR product of the central primer pair (MP5+MP6) is sequenced by Sanger sequencing to confirm faithful integration. **C.** Segregation analysis by PCR of a transgenic spirulina strain (SP607; expresses a GST-tag) over several months after transformation. Spirulina was transformed on day 0 with donor DNA containing an antibiotic marker (*aadA*) and cultured under streptomycin selection. PCR products of the full transgene region (primers MP5 and MP6) were amplified from genomic DNA samples collected at the indicated timepoints. The extent of segregation was assessed by loss of the wild-type loci band (SP3). Complete segregation was observed by day 72. **D.** Segregation analysis of a transgenic strain (SP79; expresses a woodchuck hepadnavirus virus like particle) cultured continuously for more than 3 years. PCR amplification was performed with primers targeting the full transgene region (primers MP5 and MP6).

Spirulina are polyploid. Segregation of the transgene to homozygosity typically occurred at 8-10 weeks after transformation under continuous antibiotic selection (Figure 1C). Clonal derivatives were then isolated by picking individual spirulina filaments under a microscope and verified by quantitative comparison to an endogenous chromosomal gene using qPCR as containing a single precisely integrated insertion per spirulina chromosome (see Methods). Long-term stability was assessed for 9 strains continuously propagated for at least 1 year (>800 cell generations). PCR and DNA sequencing showed all strains to be genetically indistinguishable from the original engineered strain. A strain expressing an exogenous vaccine antigen has been genetically stable during 3 years of continuous propagation (Figure 1D).

### Induction of spirulina competence

We determined that transformation competence could be induced in spirulina by co-cultivation with specific companion microorganisms. Spirulina UTEX LB1926 obtained from the UTEX algae culture collection was not axenic. Both direct microscopic examination and seeding of the culture medium onto solid LB+SOT plates revealed the presence of contaminating microorganisms. Axenic strains of spirulina UTEX LB126 were generated using micromanipulation to pick single filaments. Eight of 11 single filaments produced axenic cultures as verified by microscopic examination and confirmed by negative results after cultivation on LB+SOT plates. Three of 11 single filament-derived strains remained non-axenic. When exposed to 2 different integrating DNA vectors, all 3 xenic filament-derived strains remained naturally transformable, whereas none of the 8 axenic cultures were transformable, suggesting that competence for transformation was associated with the presence of other microorganisms.

Microorganisms found in the original xenic strain of UTEX LB1926 were streaked to single colonies. Twelve isolated colonies were individually co-cultured in liquid medium with an axenic UTEX LB1926 strain. The axenic spirulina strain remained non-transformable, but the spirulina in all 12 co-cultures became competent for natural transformation. Microorganisms that induced competence in these cultures were identified by sequencing 23S and 16S chromosomal DNA as belonging to the genera sphingomonas and microcella. Similar results were obtained for NIES-39. Therefore, we used xenic strains for genetic transformation and derived axenic variants for subsequent protein production.

### A markerless method for engineering of spirulina

The site of chromosomal integration was dictated by the homology arms flanking the transforming genes. A construct containing a single homology arm was ineffective at transformation; this indicated that integration occurred primarily by double crossover events, as is typical for natural transformation^28^. The homology arms were 1–1.5 kb in size, though efficient integration was observed with a 118 bp homology arm. To date we have evaluated 11 sites of integration, of which 6 were shown to accommodate an insertion with no apparent deficit in a standard growth regimen.

One selected insertion site corresponded to NCBI reference sequence: WP_006618409.1. A BLAST search of this gene yielded homologs from other bacteria that were annotated as kanamycin aminoglycoside acetyltransferase genes. We found that it conferred kanamycin resistance in *E. coli*, and that deletion of this locus rendered spirulina kanamycin sensitive. Kanamycin resistance encoded by this natural spirulina gene presented an opportunity for development of a markerless genetic engineering strategy. The first step used homologous recombination to precisely replace this gene with an exogenous gene (*aadA*) encoding resistance to streptomycin. The resulting strain, SP205, was kanamycin sensitive and streptomycin resistant, but otherwise identical to the parental strain. In the second step the natural spirulina kanamycin-resistance gene was joined to a gene of interest and precisely re-introduced into its original location in the genome of SP205 by replacement of the exogenous *aadA* gene. The resulting strain had kanamycin resistance restored, expressed the gene of interest, and again was streptomycin sensitive. This strain contained the gene of interest integrated into the spirulina chromosome at a pre-selected, defined location and contained no other exogenous genetic information (Figure 2).

**Figure 2:**
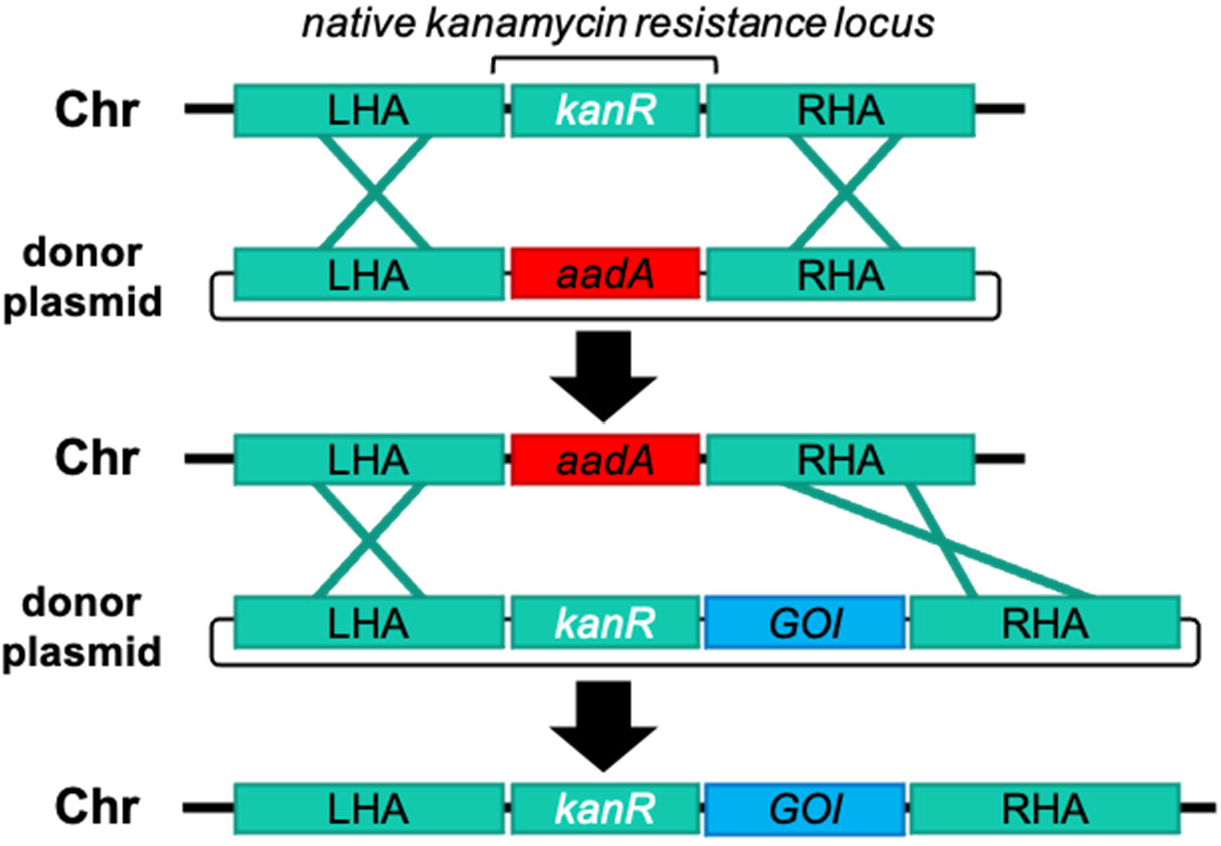
Strategy of markerless transgene integration. First, spirulina is transformed with an integrating vector with homology arms (LHA and RHA) targeting a native kanamycin-resistance gene (*kmR*) in the genome (Chr). A non-native marker conferring streptomycin resistance (*aadA*) is transformed as a donor plasmid and used to replace the native antibiotic resistance gene, making the progeny strain kanamycin sensitive. A fully segregated strain containing the *aadA* marker is then transformed with a donor plasmid that restores the native *kmR* locus sequence while introducing simultaneously introducing a recombinant transgene cassette for a GOI. The final construct contains no non-native antibiotic resistance genes and is resistant to kanamycin and sensitive to streptomycin.

To date we have demonstrated introduction of a variety of exogenous DNA vectors into the spirulina genome, including single genes, genes in tandem, and operons, as well as sequential engineering of different insertion sites has been demonstrated. The largest transforming DNA cassette introduced was 5.3 Kb and contained a 7-gene C-phycocyanin operon from a thermophilic cyanobacterium. We demonstrated intracellular expression of a diverse group of exogenous proteins, including bioactive peptides, antibodies, protein pigments, and enzymes. Self-assembling nanoparticles derived from bacteriophage or viral capsid proteins for multivalent display of vaccine antigens have also proven readily expressible. The amount of an exogenous protein expressed using a strong, constitutive promoter derived from the C-phycocyanin locus (Pcpc600) was as much as 20% of total soluble protein.

### Expression of camelid single-chain antibody fragments in spirulina

Unlike human antibodies, antigen-binding domains (VHHs) derived from camelid single-chain antibodies are ideal for expression in prokaryotes, like spirulina, because neither intracellular formation of disulfide bonds nor specific glycosylation is required for synthesis of the bioactive protein^29^. VHHs were constitutively expressed in spirulina from the strong promoter Pcpc600 in various formats, including monomers, dimers^30^, trimers^31^ and heptamers^32^ (Supplemental Figure S2). Monomeric VHHs were typically expressed as a fusion protein with a solubility-enhancing chaperone, such as the *E. coli* maltose-binding protein (MBP). Multimers were constructed using scaffolding strategies, and routinely demonstrated sub-nanomolar Kd values (Figure 3). Expression levels varied among VHHs, ranging from 0.5% to 20% of soluble protein.

**Figure 3:**
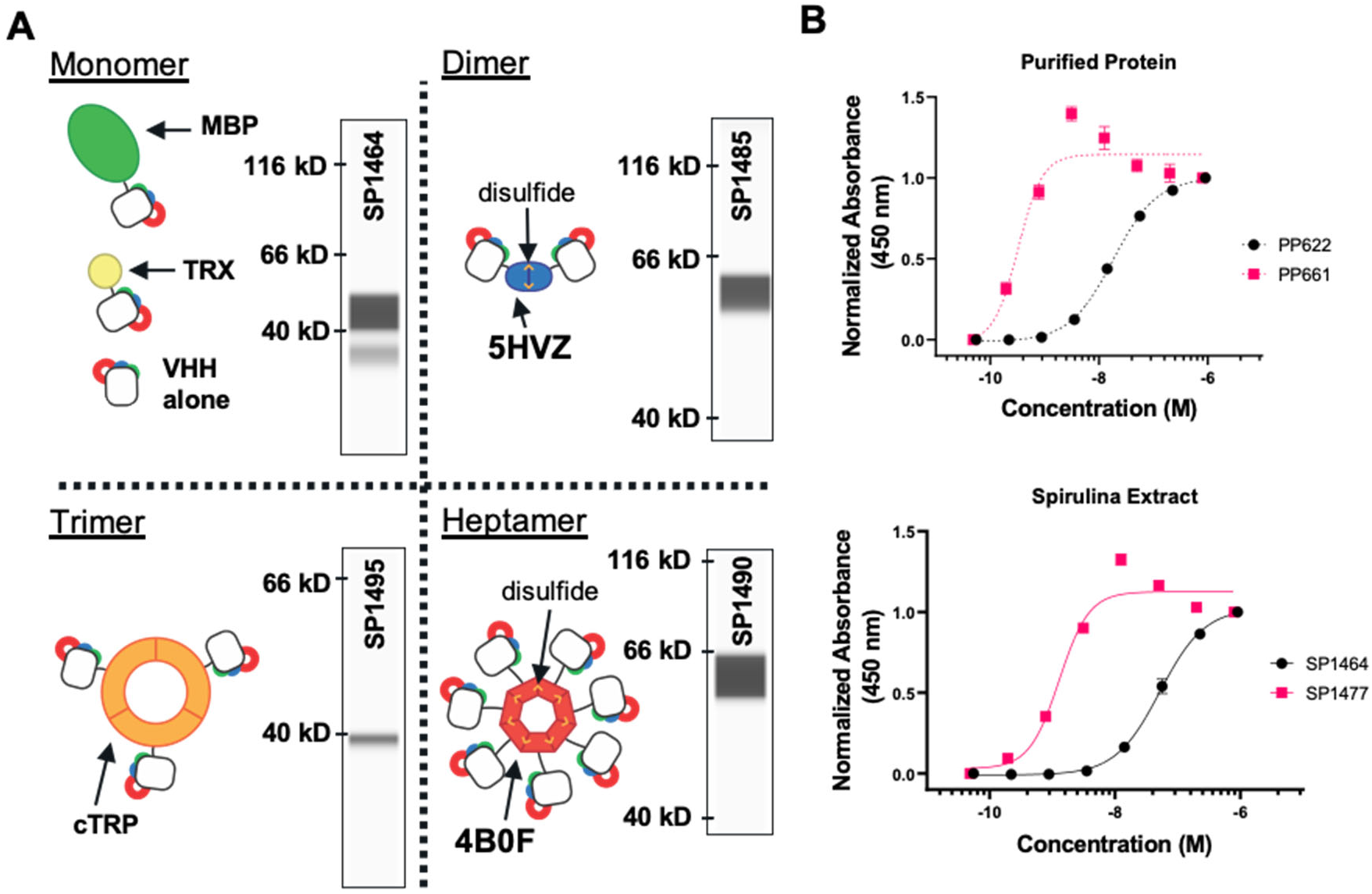
VHH scaffolding strategies. **A.** Cartoons of multimeric scaffolds with sample expression data for VHHs in spirulina. Monomeric (maltose-binding protein (MBP) and thioredoxin(TRX)), dimeric (5HVZ), trimeric (cTRP), and heptameric (4B0F) scaffolding proteins have been used to multimerize VHHs expressed in spirulina. Inter-subunit disulfides confer additional stability to the dimeric and heptameric scaffolds; these forms were commonly expressed with an MBP tag to improve solubilization. Inset capillary electrophoresis immunoassay (CEIA) blots for each scaffold demonstrate spirulina expression of a SARS-CoV2 RBD-binding VHH^68^ fused to the indicated scaffolding protein. All VHH fusions were observed at the appropriate size. **B.** Increase in binding activity by dimerization of VHHs as measured by ELISA with purified VHH (top) and spirulina extract (bottom). *E. coli*-expressed and purified monomeric (PP622) and dimeric (PP661) forms of the RBD-binding VHH were assayed with RBD and compared with the binding activity of identical proteins present in spirulina extracts (SP1464 and SP1477 respectively). The concentration of VHH in the spirulina extracts was determined by CEIA. EC50s of 18.3 nM and 52.3 nM were measured for the monomeric VHH in purified (PP622) and extract (SP1464) samples respectively, while EC50s for the dimeric VHH were 0.32 nM and 1.29 nM for the purified (PP661) and extract forms (SP1477).

Because they are easily expressed in prokaryotes, VHHs can be rapidly isolated from high diversity, naive phage-display libraries^33^. These typically have mid-nM affinities for their antigen targets, and therefore further mutagenesis may be required to achieve the higher affinities required for therapeutic efficacy. Expression of VHHs as high-avidity multimers, as described here, bypasses this requirement and therefore can accelerate product development.

### Spirulina strains expressing an anti-campylobacter VHH

The VHH FlagV6 has been reported to bind the flagellin (FlaA), a subunit of *C. jejuni* flagella with a K_D_ of 25 nM^34^. The binding site was mapped to the D3 domain of FlaA by phage display of peptides tiled across the entire FlaA protein (Figure 4A). Spirulina strain SP526 used Pcpc600 to drive expression of a monovalent fusion polypeptide of FlagV6 VHH and MBP. Two alterations were introduced into the N-terminal portion of the framework region of FlagV6 to confer increased resistance to chymotrypsin (K3Q and E5V)^35^. This anti-campylobacter protein was designated aa682. (SP1182 a markerless version of SP526 was subsequently constructed for clinical testing.) Following segregation to homozygosity, single filaments were isolated and an axenic strain was derived. Constitutively expressed aa682 protein amounts were determined by capillary electrophoresis immunoassay (CEIA) to be approximately 3% of biomass (Figure 4A). Similarly to the reported affinity of FlagV6, aa682 was found to bind to a recombinant FlaA construct with a >*K*_*D*_ of 53 nM (Figure 4B). The VHH-binding site was mapped to the D3 domain of FlaA by phage display of peptides tiled across the surface-accessible region of FlaA (see Methods).

**Figure 4:**
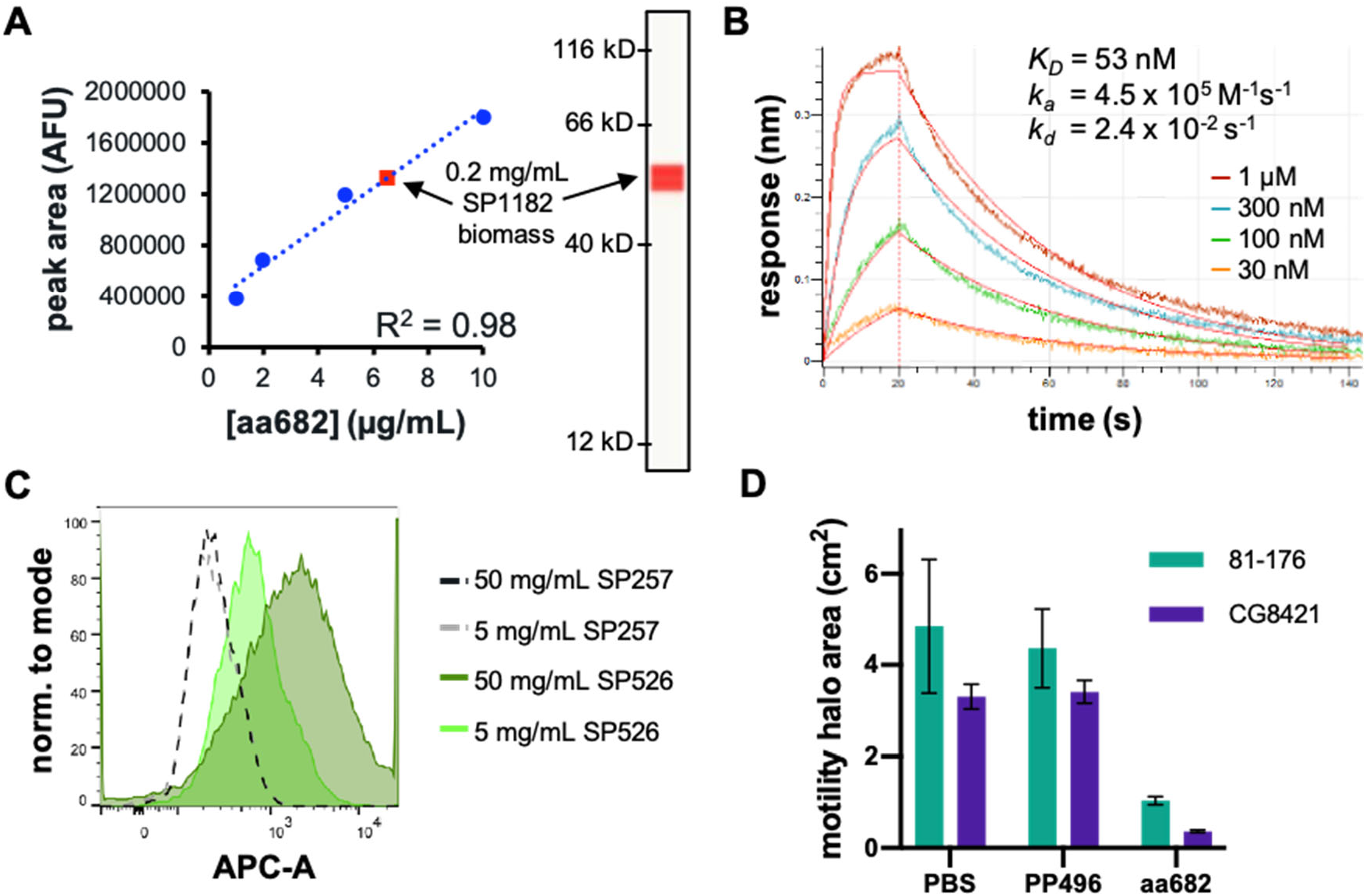
Characterization of spirulina-expressed anti-campylobacter VHH. **A.** CEIA quantification of aa682 in SP1182. A standard curve of purified aa682 protein measured on a Jess system by anti-His-Tag detection using a fluorescent secondary detection antibody in the IR channel is shown. Clarified lysate from spray-dried SP1182 was loaded at a concentration of 0.2 mg biomass/ml. A single peak was observed at the correct molecular weight of 54.8 kD and soluble aa682 was measured at ~3% of total dried biomass based using the standard curve. **B.** Binding kinetics of spirulina-expressed aa682 with recombinant flaA measured by BLI. Streptavidin-coated biosensors were loaded with biotinylated flaA and association and dissociation were measured with the indicated concentrations of aa682. Curve fitting was performed using a 1:1 binding model. **C.** Binding of VHH to intact *C. jejuni*. Soluble extracts from spray dried spirulina biomass containing an irrelevant VHH (SP257) or an analog of aa682 (SP526) were incubated with *C. jejuni* 81-176 and stained with a fluorescent anti-His-Tag antibody. Fluorescence was measured by flow cytometry. **D.** Motility inhibition of *C. jejuni* motility by aa682. Two strains of *C. jejuni* (81-176 and CG8421) were grown on soft agar plates in the presence of aa682 or an irrelevant VHH control (PP496). Samples were prepared in triplicate and motility halos were measured either 40 h (81-176) or 66 h (CG8421) after plating. Motility halo area was calculated from the halo diameter. Data represented as mean ± SD.

The D3 domain of FlaA protrudes from the axis of the flagellum, and should be surface-accessible for VHH binding^36^. Specific binding of aa682 to intact *C. jejuni* flagella was demonstrated by flow cytometry. Aqueous extracts of a spirulina strain (SP526) expressing an analog of aa682 were incubated with a pure culture of *C. jejuni* 81-176 (10^7^ colony forming units (CFU)/ml) and then stained with anti-His-Tag antibody to detect the binding of the VHHs to the pathogen. Binding to *C. jejuni* was compared to an extract from a spirulina strain (SP257) that expressed an irrelevant VHH (Figure 4C).

The major flagellin protein FlaA is required for motility, and motility is required for virulence *in vivo*^37^. Binding of VHHs to FlaA has previously been shown to prevent campylobacter motility *in vitro*^34^. Purified aa682 blocked campylobacter motility on agar plates of 2 different strains of *C. jejuni* (Figure 4D) and was therefore predicted to inhibit campylobacter pathogenesis *in vivo*.

### Preventing campylobacter disease *in vivo*

Two independent mouse models were used to test whether orally delivered spirulina biomass containing an anti-campylobacter VHH could prevent enteric campylobacter infection. In the first^38^, mice were rendered susceptible to campylobacter infection by an antibiotic pre-treatment regimen, and then challenged on day 0 with 10^6^ CFU of *C. jejuni* 81-176 by oral gavage. Spirulina biomass used for treatment and control groups was cultured in mid-scale bioreactors, washed, and spray dried. Untreated controls, or controls treated with either spirulina biomass containing no recombinant protein (SP227; engineered for phycocyanin production), or spirulina biomass containing an irrelevant VHH, were compared to mice given SP526 by oral gavage prior to campylobacter challenge, and again on days 1 and 2. Treatment with SP526 reduced campylobacter fecal shedding by 3 to 4 orders of magnitude, and significantly reduced two biomarkers of intestinal inflammation, lipocalin-2 (LCN-2) and myeloperoxidase (MPO) (Figure 5A, 5B). It was observed that all of the mice in the control groups, but none of the mice treated with SP526, suffered from diarrhea following campylobacter infection. The severity of this overt clinical disease was not quantified.

**Figure 5:**
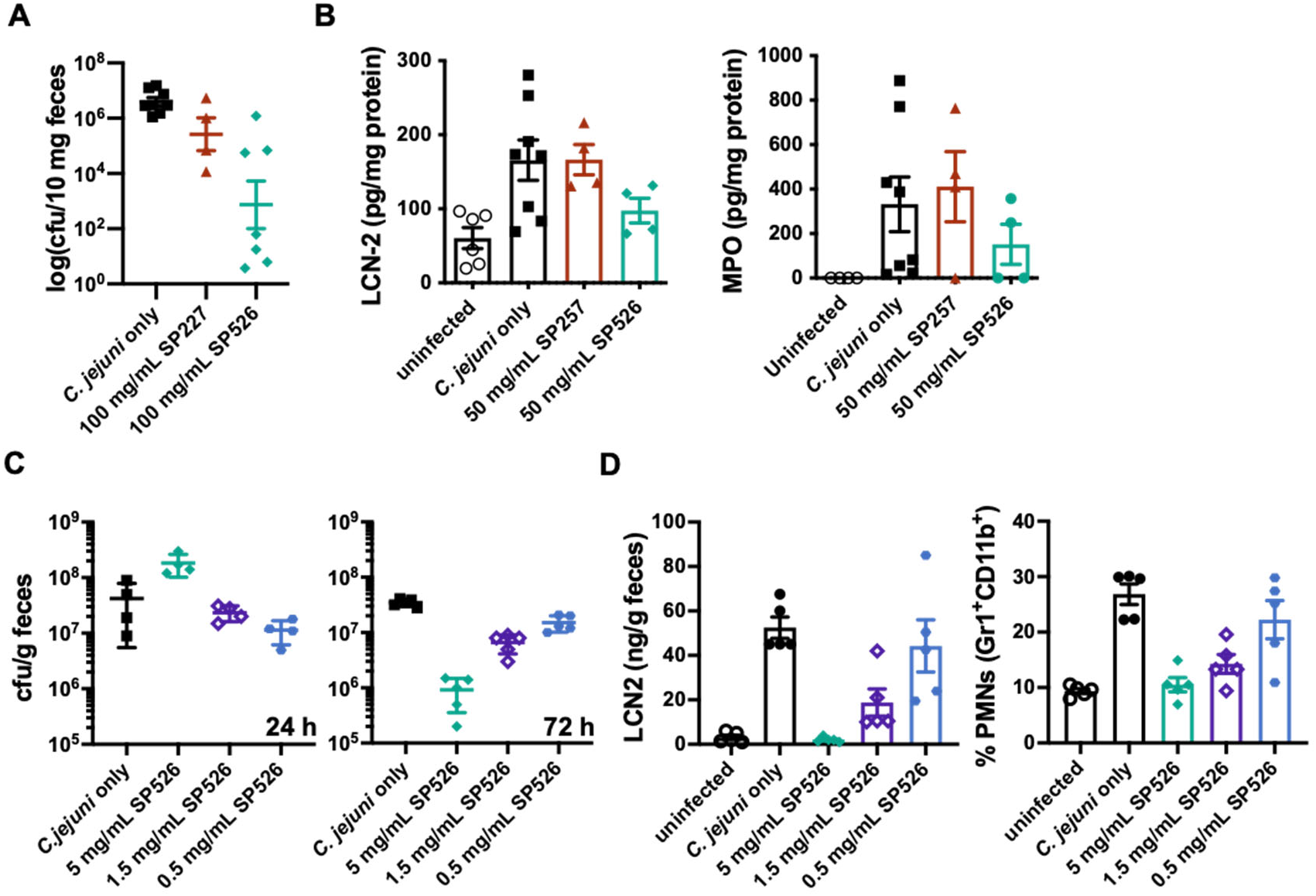
Prevention of *C. jejuni* infection in mice. **A.** Shedding of *C. jejuni* in a mouse model of infection. In a pilot experiment, mice received a daily 200 μL dose spirulina biomass resuspensions or vehicle between days −1 and +3 relative to challenge (5 total doses). Spirulina strain SP227 expressed no VHH and SP526 expressed an analog of aa682. Bacterial shedding in stool was measured 7 days after challenge. **B.** Biomarkers of inflammation (LCN-2 and MYO) were measured in stool 11 days after infection. Mice received a daily 200 μL dose of spirulina biomass resuspensions or vehicle on days −1, 0, +1 relative to challenge. Spirulina strain SP257 expressed an irrelevant VHH. **C.** Bacterial shedding after treatment with a single dose of SP526. Mice received a single 400 μL dose of spirulina resuspension or vehicle 1.5 h before challenge with *C. jejuni*. Bacterial shedding in stool was measured 24 and 72 h after challenge. **D.** Markers of inflammation (LCN2 and PMNs) were measured 72 h after challenge and treatment with a single dose of SP526. All data represented as mean ± SEM.

A second model also used an antibiotic preconditioning regimen but a challenge dose of 10^8^ CFU of *C. jejuni* 81-176. Dose-ranging experiments showed that a single prophylactic dose of 2 mg of dry SP526 biomass administered by oral gavage was sufficient to prevent campylobacteria disease, as measured by stool lipocalin-2 and myeloid cell infiltration (PMNs) of the cecum. Furthermore, this single dose accelerated campylobacter expulsion from the gut in the first 24 hours after challenge, which was accompanied by reduced campylobacter shedding at 72 hours after challenge (Figure 5C, 5D).

### Large-scale continuous growth

Open pond systems are typically used to cultivate spirulina at commercial scale for production of food, feed, and pigments, but uncontrolled exposure to environmental contaminants make these challenging for the manufacture of biopharmaceuticals under FDA cGMP. Therefore, we developed and validated an indoor, pH-controlled, air-mixed photobioreactor platform built around a modular 160L-2,000 L vertical flat-panel reactor that is scalable to commercial size suitable for the manufacturing of biopharmaceuticals.

We found that major advantages of this platform are the exceptionally low cost of large-scale growth and downstream processing. A key factor contributing to the low upstream production cost was that spirulina thrive under extreme conditions (pH>10 and high total salinity) that severely limit the growth of adventitious organisms. In addition, spirulina, being photoautotrophic, have simple nutritional requirements; no carbon-based source of energy (i.e. sugar feedstock) is required. Together these features allowed the use of unsealed reactors under sanitary, but not aseptic, conditions. Total microbial counts in the final product were within specified limits, and absence of pathogenic bacterial confirmed regularly. Utilizing sodium nitrate allowed for high nitrogen levels in the formulated media without the toxicity that could result from the use of ammonia, and a high initial nitrogen level obviated the need for monitoring or re-feeding of nitrogen during growth. Single-use polyethylene bags contained the spirulina culture, further reducing what is typically one of the biggest cost components in any biopharmaceutical process: sterilization downtime.

The energy cost for LED illumination was the major component of production cost (Figure 6A). The reactors were illuminated from both sides of the culture using off-the shelf commercial full-spectrum LEDs with adjustable intensity. The complex relationship between the capital and operational costs of biomass growth were evaluated as a function of light intensity, and an optimum that achieved the greatest productivity per unit energy cost was identified (Figure 6B).

**Figure 6:**
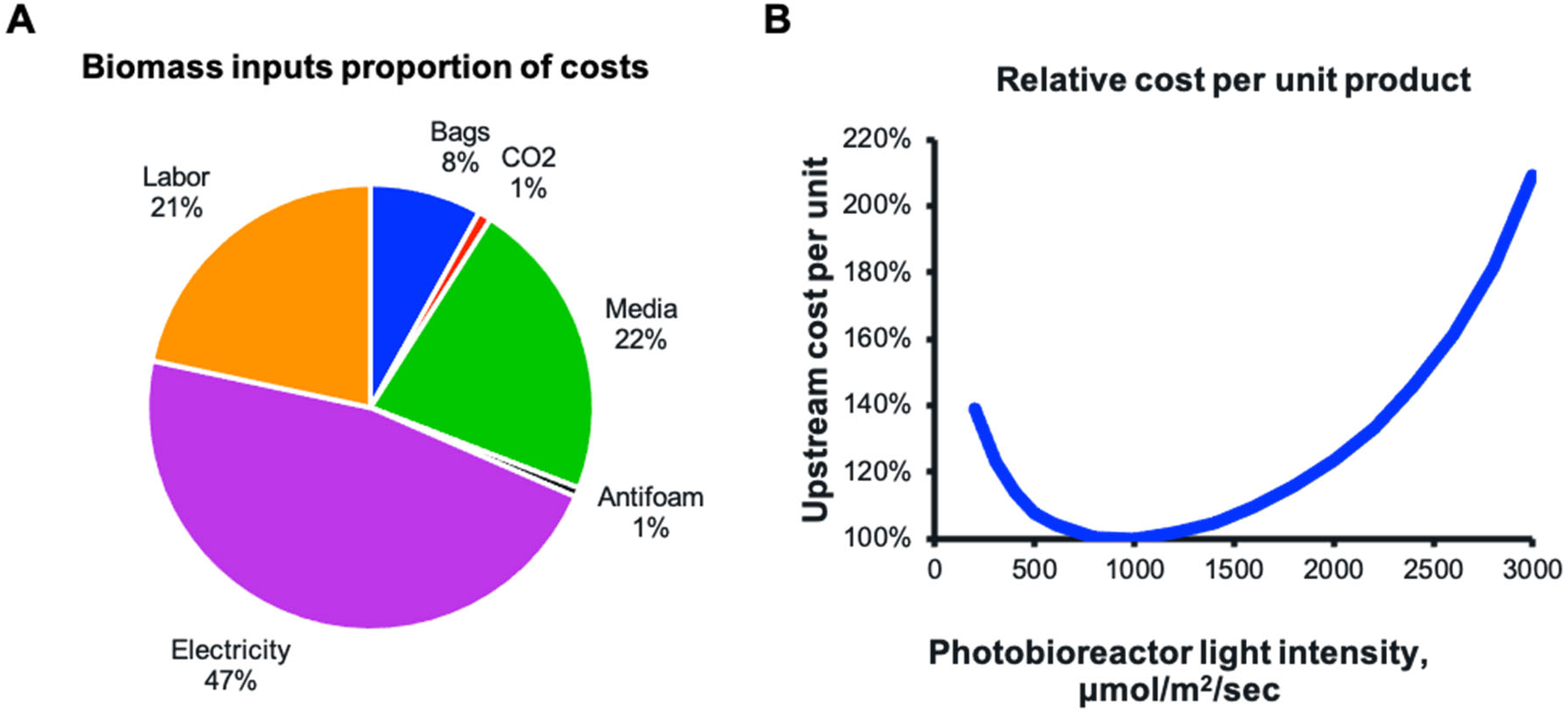
Cost optimization. **A.** Cost components of cGMP biomass production. **B.** Spirulina productivity is a function of light intensity and is empirically determined in the described system with SP1182 as the production organism and with current operating parameters. Cost per unit biomass includes labor, capitalized cost of operating lighting system (varies by light intensity), and capitalized costs of other upstream components (independent of light intensity). Minimal cost per unit biomass was achieved at a light intensity of approximately 100 μmol/m^2^/sec.

Cultures were continuously maintained for sequential 1-week growth cycles. On a weekly basis, cell densities reached approximately 4 grams/liter, at which point the biomass was harvested by pumping over a series of stainless-steel screens to concentrate and rinse the slurry. A portion of the slurry was used to re-inoculate the reactors, and the remainder was processed into drug product.

### Simple downstream processing

The concentrated spirulina slurry was rinsed with a dilute trehalose solution, and then spray dried. A large parameter space was evaluated at laboratory scale for efficiency of drying, moisture content, and retention of antibody activity. Conditions were identified (Supplemental Figure S3) in which suitable system efficiency was achieved while maintaining greater than 90% of antibody activity. The process was translated to a larger scale (5 kg/hr) spray dryer equipped with a centrifugal atomization system. With only minor optimization of system throughput, inlet and outlet temperatures and air flow rates, equivalent dryer performance at a ~20x single step scale-up was achieved.

Once dried, the powder was collected and sealed in light- and moisture-proof packaging; antibody activity was retained at room temperature for at least 1 year (Figure 7). Packaging the powder into standard vegetarian capsules was the final downstream process. Because of product stability, distribution can be cold-chain independent. The system was third party audited and found compliant with ISO 22000 Food Safety Management System standards and subsequently required minimal upgrades to satisfy pharmaceutical cGMP requirements.

**Figure 7:**
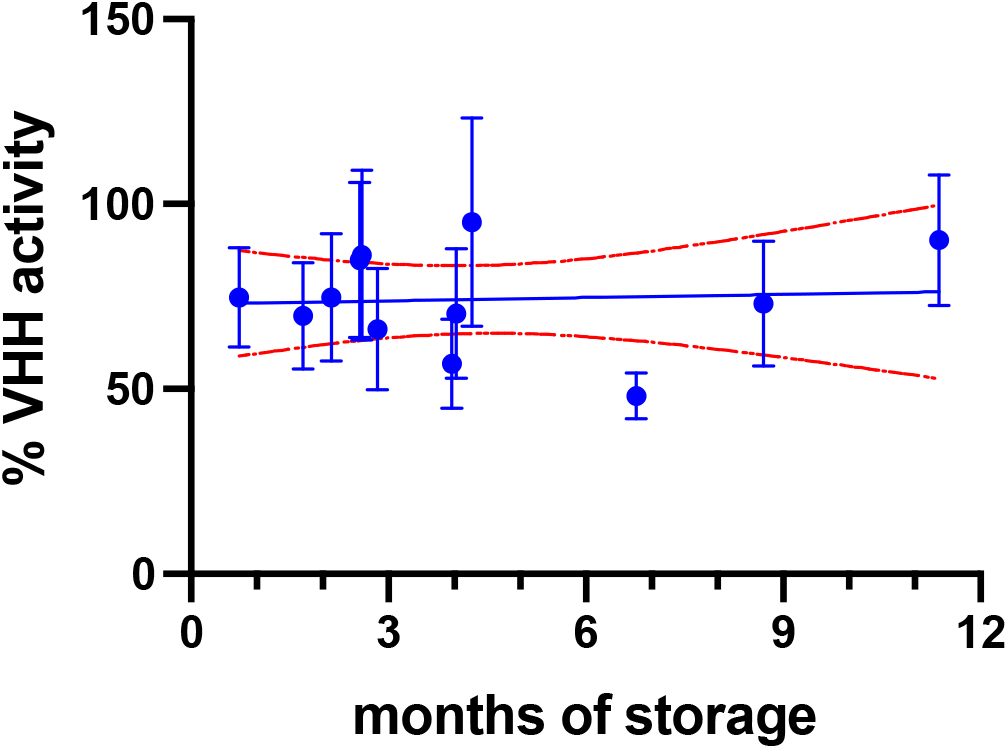
Stability testing of SP1182 binding activity. Batches of spray-dried SP1182 were stored at room temperature for the indicated amount of time and binding activity was assessed by ELISA. Samples were tested in duplicate. Binding of purified aa682 to recombinant flaA was used to generate a standard curve by linear regression. The standard curve was used to calculate the concentration of active aa682 in SP1182 lysates based on ELISA binding activity. VHH activity was normalized to 100% assuming an expression level of 3% aa682 per unit of biomass. Each point represents a different batch of biomass, and data are presented as mean ± SD. Red dotted lines indicate upper and lower 95% CI of linear regression analysis of samples.

### Targeted delivery

A significant challenge associated with direct delivery of protein therapeutics to the gastrointestinal tract is protease digestion. Upon ingestion, biologics are initially subjected to the low pH and high pepsin gastric environment, so we evaluated whether bioencapsulation of therapeutic proteins within dry spirulina biomass would provide protection. As expected, purified aa682 was fully degraded within 2 minutes of incubation in simulated gastric conditions (Figure 8A). However, when delivered within dry spirulina biomass more than 70% of aa682 remained intact after 2 hours of incubation in the same gastric conditions (Figure 8B). Transition of biomass to the higher pH of a simulated duodenal environment was then sufficient to extract more than 90% of the encapsulated aa682 from spirulina within 60 minutes (Figure 8C). When assayed by ELISA, the binding activity of the extracted aa682 was unaffected by the initial biomass incubation in the simulated gastric environment (Figure 8D).

**Figure 8.**
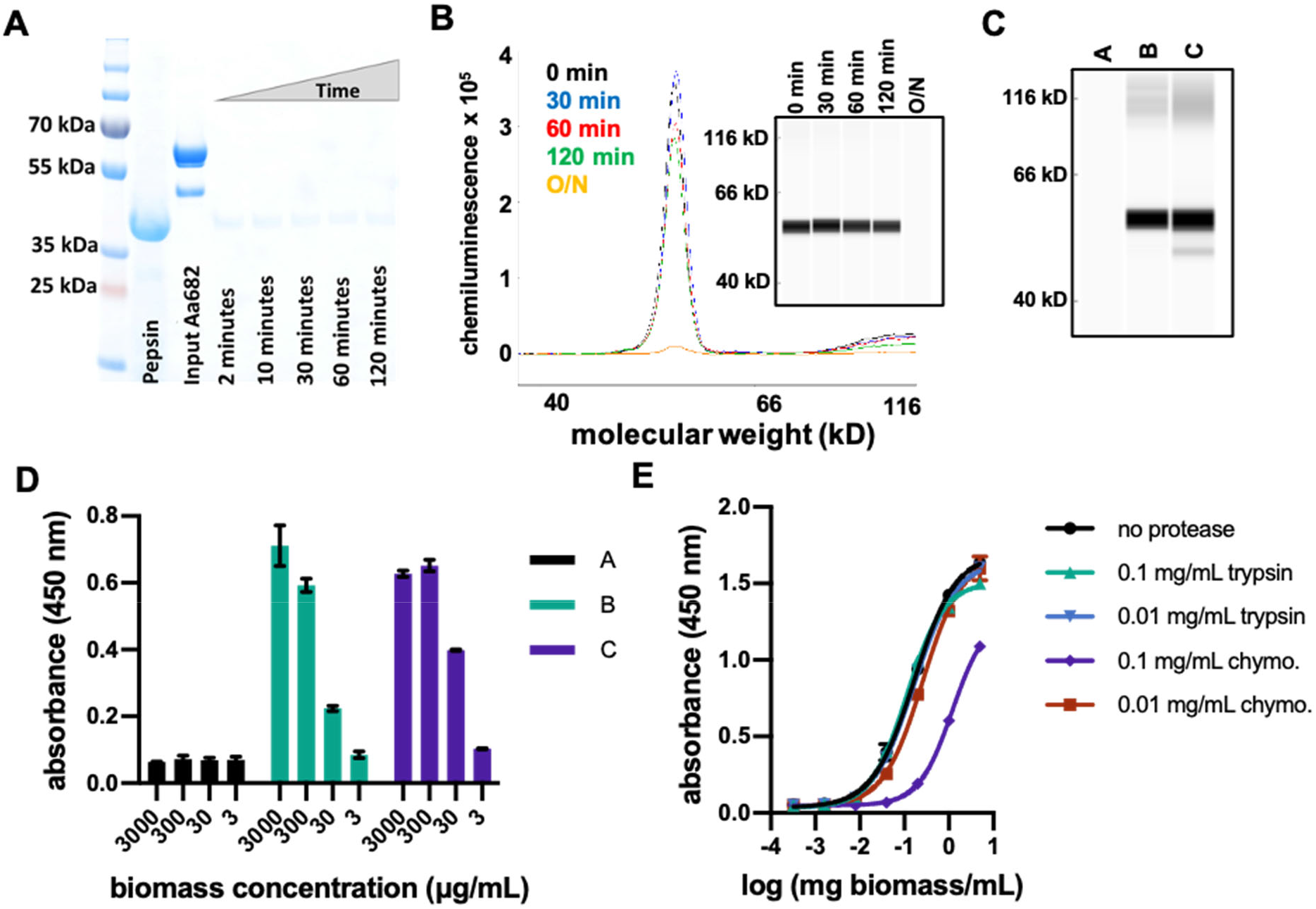
*In vitro* analyses of protease resistance of aa682 and SP1182 biomass **A.** SDS-PAGE analysis of purified aa682 incubated with simulated gastric fluid supplemented with 2000 U/ml pepsin. Pepsin band is indicated by loading 20 μg of pepsin. Digestion of aa682 was quenched at intervals ranging from 2 to 120 min. This data is representative of 2 independent experiments. **B.** CEIA of pepsin-digested spirulina biomass resuspension. Dried spirulina biomass of SP1182 was resuspended in simulated gastric buffer and incubated with pepsin for 0 to 120 min or overnight. Whole biomass samples were denatured and analyzed on a Jess system. Recombinant aa682 was detected with an anti-His-Tag antibody and the data are representative of 4 independent experiments. **C.** Immunoassay analysis of spirulina biomass resuspended in low pH gastric simulating buffer conditions. Dried spirulina biomass of SP1182 was resuspended in simulated gastric buffer (pH 3.0, no pepsin) and the presence of aa682 was analyzed in soluble buffer (A). The soluble fraction of biomass subsequently pelleted and resuspended at a higher pH (B) was compared to aa682 extraction by direct resuspension in a high pH bicarbonate buffer (C). The data are representative of 2 independent experiments. **D.** ELISA-based antigen-binding analysis of spirulina extracts from biomass resuspended in gastric-simulated buffer condition. Soluble protein extracts of (A), (B), and (C) were prepared as in figure 8C were assayed for antigen binding to recombinant flaA. Samples (B) and (C) yielded approximate EC50 binding values of 85.8 μg/ml and 29.2 μg/ml of biomass, respectively. Data presented are averages of 2 replicates. **E.** ELISA binding activity of aa682 after *in vitro* protease digestion with intestinal proteases. Lysates from SP1182 were incubated with 0.1 or 0.01 mg/ml of trypsin or chymotrypsin (chymo.) for 1 hr. After protease activity was neutralized, aa682 binding activity to recombinant flaA was measured by ELISA. Bound VHH was detected with an anti-VHH antibody cocktail. Samples were assayed in duplicate.

*In vivo* efficacy is also dependent upon the sensitivities of therapeutic proteins to proteases of the small intestine, especially trypsin and chymotrypsin. Empirical analysis is necessary to identify VHHs with appropriate metabolism-resistant properties, which range from highly sensitive to almost complete resistance to intestinal proteases. When soluble extract from SP1182 was treated with proteases, aa682 was resistant to the constitutive intestinal levels of both trypsin and chymotrypsin. Some sensitivity of aa682 to chymotrypsin, but not trypsin, was evident when the SP1182 extract was exposed to the high induced protease levels that are present immediately after ingestion of a meal^39^ (Figure 8E).

### First-in-human clinical trial

Spirulina strain SP1182 was cultured in large-scale bioreactors under cGMP conditions (see Methods) and used to formulate the drug product LMN-101. To assess the safety and tolerability of LMN-101, a Phase 1 clinical trial was designed and conducted. Eligible, healthy volunteers aged 18-50 were enrolled following informed consent. The study was performed in accordance with ICH guidelines and in compliance with all applicable laws and regulations.

A total of 20 subjects were randomized to active or placebo treatment as described in the following sequence:

- 3000 mg of LMN-101, administered orally as 6 500-mg capsules as a single dose (2 subjects)
- 300 mg of LMN-101, administered orally as a single 300-mg capsule 3 times daily for 28 days (4 subjects) or identical-appearing placebo capsule (2 subjects).
- 1000 mg of LMN-101, administered orally as 2 500-mg capsules 3 times daily for 28 days (4 subjects) or identical-appearing placebo capsules (2 subjects).
- 3000 mg of LMN-101, administered orally as 6 500-mg capsules 3 times daily for 28 days (4 subjects) or identical-appearing placebo capsules (2 subjects). 1 additional subject in this cohort who dropped off study was replaced.

This Phase 1 trial demonstrated LMN-101 was safe and well tolerated at the doses tested. There were no statistically significant differences between the active and placebo groups. There were no significant adverse events or significant laboratory abnormalities reported during or following the trial. Rates of adverse events, deemed possibly related, were similar between the two groups. Reported adverse events were mild and consisted of nausea, abdominal pain, diarrhea, gastroesophageal reflux, constipation, pharyngitis, and delayed menstruation. Laboratory evaluation demonstrated there was no significant VHH absorption.

## DISCUSSION

Advances in biotechnology have often followed the introduction of new platform organisms to enable production of biopharmaceuticals with new characteristics and which unlock treatments for diseases unresolved by older technologies. Spirulina’s advantages as a biotechnology platform include not only low cost manufacturing and cold-chain independent distribution, but also its intrinsic safety which can simplify and de-risk the drug development process. Spirulina holds FDA Generally Recognized as Safe (GRAS) status due to its long history of safe human consumption^40^. Furthermore, its safety and toxicology profile has been positively reviewed by the FDA, and numerous clinical trials have established its safety for consumption both adult^41–44^ and pediatric populations^45^.

Targeted delivery of any therapeutic agent to its site of action is a key consideration for efficacy, and is another advantage of the spirulina platform for treatment of enteric diseases. Typically, therapeutic agents are delivered by oral or injection pathways and are then subjected to absorption, distribution, metabolism, and elimination effects that reduce the amount of active product present at the site of action, thus reducing efficacy. These challenges are exacerbated by the risk of toxicity at higher dose levels for all injected biopharmaceuticals. Referred to collectively as “ADMET”, these issues are often reasons for clinical failure and a major driver of the skyrocketing cost of new drug development^46^.

By contrast, delivering a therapeutic biomolecule that is not absorbed by the intestine within a safe food substance reduces these concerns. Considerations of distribution are inconsequential as the product is delivered directly to the site of disease. Elimination, through the function of the GI tract, is linear and predictable; most importantly, it does not reduce the amount of active agent at the site of action, although transit time can affect the duration of action. Digestion of the therapeutic within the GI tract is a challenge that must be addressed, but is outweighed by lower ADMET costs and risks. The therapeutic approach therefore compares favorably with systemically dosed therapeutics for common GI diseases such as bacterial enteritis^47^ and inflammatory bowel disease^48^ which have well-documented side effects.

### Implications for campylobacter disease

Enteric infectious diseases remain prevalent today, including those caused by *C. jejuni,* enterotoxigenic *E. coli,* shigella and *Clostridioides difficile*. Moreover, some of these are designated high-priority antimicrobial resistance threats by the US Centers for Disease Control and Prevention, which has added urgency to the search for new therapeutic tools. Diarrhea still accounts for nearly 10% of the 7.6 million deaths in children under the age of 5 annually^49^, and campylobacter is among the most common causes of bacterial gastroenteritis worldwide^50–52,53,54^. It is particularly hazardous to infants in the developing world and is a leading cause of infant mortality^53^. The global health burden of campylobacter diarrhea is estimated to be 88 million cases in children under 5 years of age, resulting in 41,000 deaths annually^55^. There are 172 million cases globally in all age groups, with 75,000 deaths annually. Morbidity and mortality are due not only to the campylobacter itself, but also to secondary gut dysfunction leading to malnutrition, growth stunting and increased susceptibility to additional infectious diseases.

The minimal effective dose of the anti-campylobacter aa682 in the mouse disease models was approximately 60 μg—a final concentration of approximately 1 micromolar if distributed over the 1 ml fluid volume of the mouse small intestine. This is approximately 30-fold greater than the K_D_ of the aa682 for campylobacter FlaA, which likely reflects the need to maintain a therapeutic level of the protein in the face of ongoing proteolysis. This can be mitigated, to some extent, by selecting lead VHH candidates with high levels of protease resistance. Nevertheless, it was possible to administer doses that exceeded even this high amount by 10-fold, both in mice and in a human Phase 1 trial. The lack of any toxicity demonstrated the therapeutic advantage afforded by this safe platform.

The spirulina platform is well suited for delivering high antibody doses orally, not only because of its safety but also because of the very low costs of growth and processing. The estimated adult human dose of LMN-101 would be 200 mg (containing 6 mg of anti-campylobacter aa682) based on the relative compartment volume of 100 ml for the human small intestine^56,57^ versus 1 ml for the mouse^58,59^. At full commercial scale, production under cGMP conditions of dry spirulina biomass containing a therapeutic protein is estimated to have a cost of just $100/kg implying a cost per dose below 5 cents. Significant further reductions in cost per dose may be achievable by increasing production efficiency, and increasing the expression, proteolytic stability, and avidity of the active therapeutic proteins.

Low cost and the lack of cold chain requirements make prophylactic administration practical, and moreover suitable for application in the developing world where cost is often an overriding issue. Unlike administration of conventional antibody-based therapeutics, the spirulina platform eliminates the need for cold chain storage and distribution. Maintaining antibody at temperatures from 4°C to, in some cases, −80°C creates enormous logistical challenges in developed countries and can be insurmountable in the developing world. In contrast, VHHs and other biologics simply processed into a dry spirulina powder are shelf stable without refrigeration for at least 1 year.

The efficacy of oral antibodies for the treatment of gastrointestinal infection was first demonstrated more than 40 years ago for the treatment of *E. coli* infection in human infants^60–62^. Additional successful human clinical trials against other GI pathogens, such as rotavirus and *C. difficile*, have been reported^63–67^. While these studies established the concept that orally delivered antibodies can be an effective therapeutic modality, most studies relied upon polyclonal antibodies derived from the milk of hyperimmunized cows, a production system that has proven difficult to scale cost effectively while maintaining FDA-grade lot-to-lot consistency.

Beyond the illustrative example of campylobacter disease, the spirulina platform is currently being used to develop oral therapeutics to prevent infections with other enteric pathogens, including enterotoxigenic *E. coli*, norovirus, *C. difficile,* and SARS-CoV-2. Other GI targets such as inflammatory and metabolic diseases, and, more speculatively, manipulation of the microbiome, should also be considered. While the advantages of the spirulina platform for oral delivery are immediately apparent, many of the same advantages should apply to other topical and mucosal surfaces. Taken together, the unique features of the platform make it practical to develop high-complexity biologic cocktails which increase the breadth of pathogen coverage and anti-pathogen potency, as well as potency against other disease targets. Cocktails comprised of 10 or more VHHs are in development with this platform, whereas therapeutics comprised of more than 2 antibodies are generally unfeasible using conventional platforms.

## ACKNOWLEDGEMENTS

This work was supported in part by funding from The Bill & Melinda Gates Foundation. For support especially during the critical formative phases of this work we thank Margaret McCormick, Karen Wolfe, Tom Todaro, Ed Schlect, and Dan Baty. We thank Charles Shoemaker, Barry Stoddard, George McDonald, Charles Sherr, Fred Cross, Bruce Kerwin, Kathryn Stein, Wilbur Chen, Dan Wattendorf, Omar Vandal, and members of the Vaccine Analytics and Formulation Center at University of Kansas for ongoing help, discussions and advice.

