## Supplementary Figures for "Expression and manufacturing of protein therapeutics in spirulina"

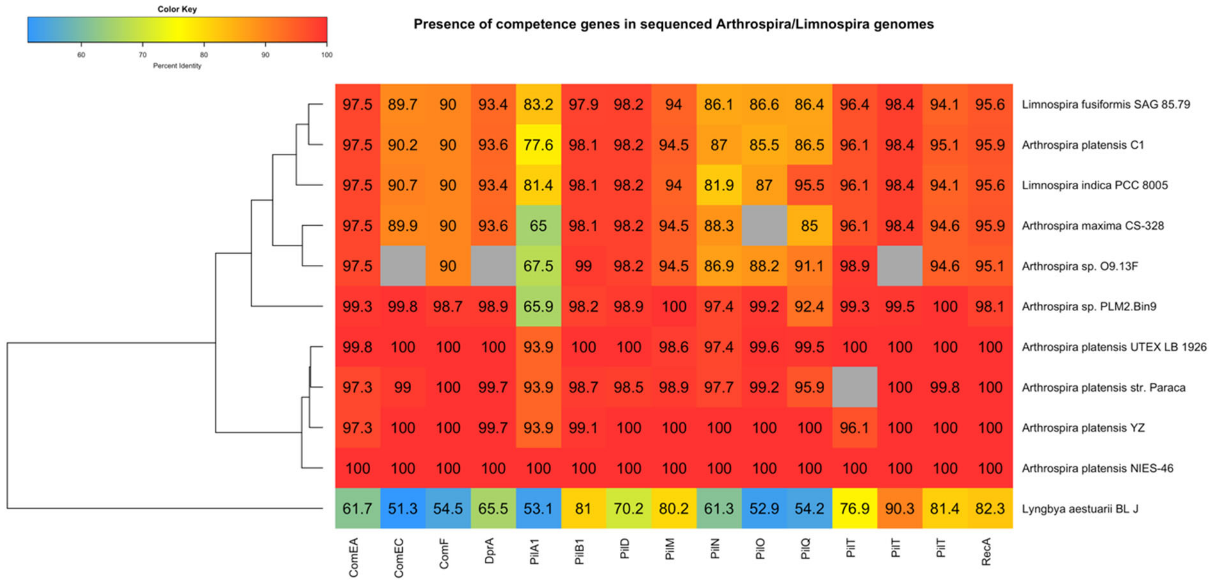


**Supplemental Figure S1:** Table of competence genes. The presence of competence genes in sequenced arthrospira/limnospira genomes was determined using reciprocal best hits against the *A. platensis* NIES-39 competence genes previously identified in Taton *et al.*, 2020. Genomes were retrieved on GenBank and BLASTp was used to identify reciprocal hits with e-values <1$^{-5}$ and query coverage >50%. Cell labels indicate the percent identity relative to the respective NIES-39 gene; cells in grey indicate that no homolog was identified*.* *Lyngbya aestuarii* BL J was included as an outgroup.

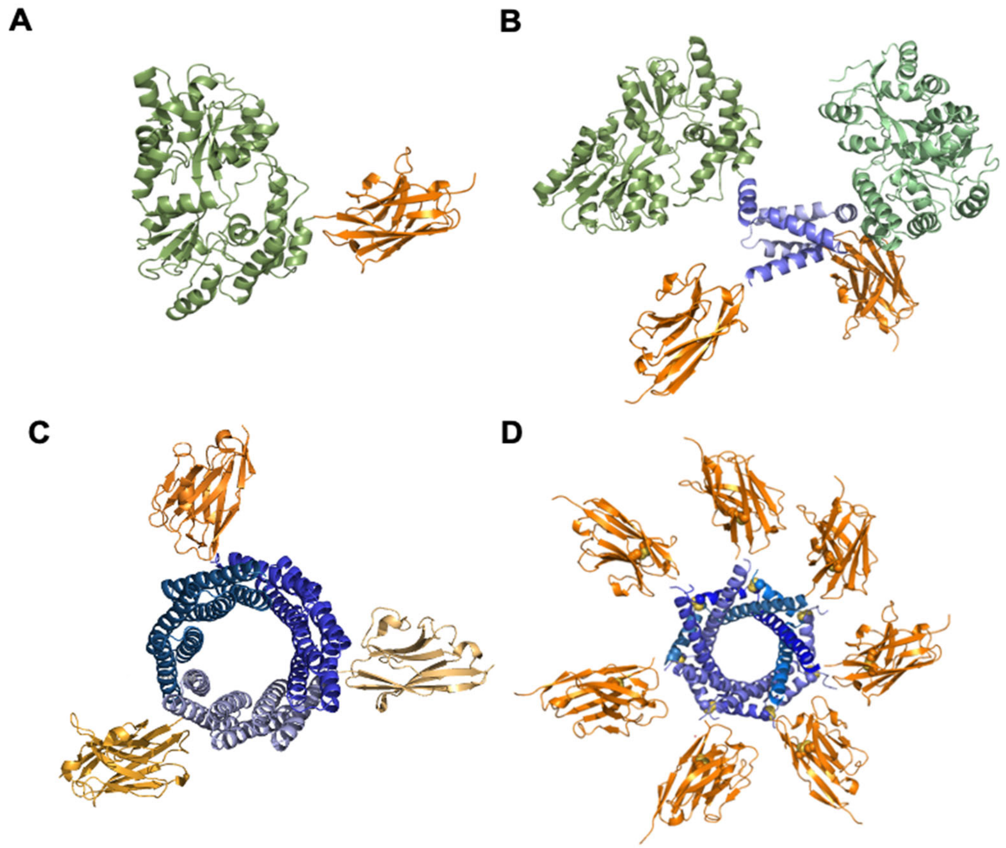


**Supplemental Figure S2.** Model representations of heterologous proteins designed for expression in spirulina. **A.** Ribbon representation of a monomeric VHH (orange; PDB ID:6WAQ)[68] with the solubility enhancer, MBP (green; PDB ID: 5M13)[71]. The mature, folded protein results in a monomeric VHH as a fusion to MBP and a C-termini 6X-his affinity tag. **B.** Ribbon representation of a VHH (orange) with a dimerization motif (blue; PDB ID: 5HVZ)[30] and the solubility enhancer, MBP (green). The mature, folded protein results in a dimeric VHH where dimerization is facilitated by the disulfide-linked dimerization motif. The single polypeptide also contains the solubility enhancer MBP and C-terminal 6X-his affinity tag. **C.** Ribbon representation of a trimeric VHH (orange). The mature, folded protein results in trimeric VHH (orange) where trimerization is facilitated by the self-assembling homotrimer t-cTRP9$X_3$ (blue; Hallinan J., *et al.* Structures and behavior of de novo designed circular tandem repeat proteins with novel repeat topologies and increased contact surfaces and thickness. In preparation.). The single polypeptide also contains a C-terminal 6X-his affinity tag. **D.** Ribbon representation of heptameric VHH (orange) with the heptamerization motif (blue; PDB ID: 4B0F)[32]. The mature, folded protein results in a heptameric VHH where heptamerization is due to intrachain disulfide bond between individual protomers. The polypeptide also contains an N-terminal solubility enhancer MBP fusion and C-terminal 6X-his affinity tag.

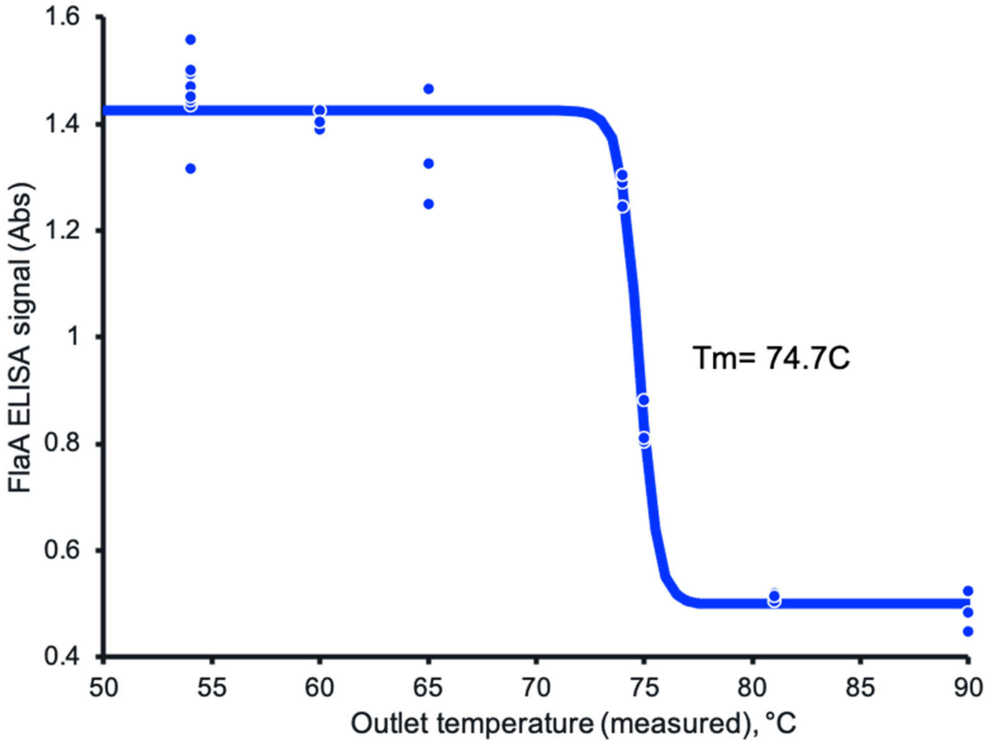


**Supplemental Figure S3:** VHH stability during spray drying. FlaA binding activity of aa682 in biomass versus drying temperature. Biomass was dried across a range of temperatures, extracted at 10 mg/ml biomass, and the extracts were diluted to a constant 0.039 mg/ml assay concentration. Binding activity of the extracts to FlaA was measured by ELISA. Binding activity was unaffected by drying temperatures <73°C.
