## Supplementary material for "Expression and manufacturing of protein therapeutics in spirulina": Materials and Methods

### METHODS AND MATERIALS:

**Small-scale spirulina culturing**
Spirulina strains were grown in liquid culture using SOT media. For antibiotic selection, media was supplemented with 70-100 µg/ml of kanamycin or 2.5-5 µg/ml streptomycin. Culture volumes ranged from 3-100 ml. In preparing strains for transformation or downstream processing, cultures were grown in Multitron incubators at 35°C, 0.5% $CO_2$, 110-150 µEi of light, and shaking at 120-270 rpm depending on culture volume. Long-term cultures were maintained by incubation in Innova incubators at 30°C, atmospheric $C0_2$, 50-110 µEi of light, and shaking at 120 rpm.

**Design of integrating vectors for spirulina transformation**
For each genomic loci targeted for integration, PCR primers with 18-20 bp overlapping sequence with a vector backbone were designed to amplify 1-1.5 kb DNA fragments from the 5'- and 3'-regions flanking the locus. These regions represented the left homology arms (LHA) and right homology arms (RHA) respectively. Gel-purified fragments were assembled with the linearized backbone vector, which contained a p15 origin and an *E. coli* ampicillin resistance marker, by Gibson assembly. Markers for antibiotic resistance in spirulina were cloned in between the 2 homology arms of the plasmid.

**Transformation of spirulina**
Spirulina cultures were grown for 3 days in Innova to reach an OD700/ml of 0.5-1. A volume of 50 ml of cells was harvested by centrifugation for 10 min at 1,600 x rcf. Cells were washed once with SOT media at room temperature. Cells were resuspended in 2 ml of SOT. A 30 µL aliquot of cells was mixed with 300 ng of plasmid DNA and incubated at room temperature for 3 h. Samples were transferred to 0.6 ml of SOT media in 14 ml round bottom tubes and incubated overnight at 25-30°C in 50-100 µEi of fluorescent light on a light rack. Each tube received 2.4 ml of SOT with appropriate antibiotics and was incubated in Multitron conditions to start the selection. For the first 20-30 days, culture medium was changed every 3 to 5 days. After 30 days, when the transformants were robustly growing, cells were diluted every 3-5 days to facilitate segregation.

**Genotyping of transformed spirulina**
Genomic DNA was prepared from spirulina cells by digestion with proteinase K. Briefly, 0.2-0.5 OD700 of cells was washed once with sterilized water. A 30 µL sample of cell pellet was mixed with 120 µL of buffer EB (10 mM Tris-Cl, pH 8.5). Proteinase K was added to the samples at a final concentration of 0.2 mg/ml. Samples were incubated at 56°C for 1 h followed by 95°C for 10 min to deactivate proteinase K. Samples were centrifuged briefly to pellet cell debris. A 1 µL sample of the supernatant was used per genotyping PCR reaction. Specific integration of the transgenic cassette was determined by separate PCRs for each homology arm. For each PCR, 1 primer annealed to a genomic sequence outside the homology arm and the other annealed to a region within the transgene. Segregation of the chromosome was assessed using a primer pair annealing to regions within the RHA and LHA. The segregation PCR yielded fragments of 2 different sizes: one from the wild-type allele and the other from the transgenic allele. Strains were considered fully segregated when no wild-type allele amplicon was observed.

To verify the sequence of the transgene, PCR was performed with the genomic DNA to amplify the fragments which includes the transgene, the homology arms, and 500 bp flanking each homology arm. PCR products were separated by electrophoresis on an agarose gel and the amplified bands were gel extracted using the Qiagen Gel Extraction kit. The purified PCR products were sequenced to verify the integrated gene and the surrounding sequences.

To exclude the possibility of cross-contamination with other strains, PCR was performed to check other loci that have been used for integration of exogenous genes. PCR of genomic DNA using locus specific primers was performed and fragment size was analyzed by agarose gel electrophoresis. DNA fragments were gel extracted and characterized by Sanger sequencing. A strain was considered free of other spirulina strains if only wild-type loci were observed.

**Transformation of barcoded integrating plasmids**
To evaluate the number of individual successful integration events per transformation, a library of DNA barcodes was transformed into spirulina and quantified by next-generation sequencing (NGS). Briefly, a 19 N barcode was cloned adjacent to an antibiotic marker (*aadA*) in a plasmid containing homology arms for integration. The barcode library was estimated to contain >8 x $10^7$ transformants. The library was transformed into strain SP3 in triplicate, following the transformation method described above, and cultured with streptomycin. Spirulina samples were collected 22 and 28 days after transformation. Genomic DNA was extracted from spirulina and used in a PCR reaction to prepare ~320 bp amplicons of the barcoded regions for NGS analysis on a MiSeq (Illumina). Sequencing reads were filtered for quality and analyzed to minimize the false positives. Counting only barcodes that were 1) present at both timepoints within a replicate, 2) unique

to each replicate, and 3) observed more than 30 times within a replicate yielded an estimated number of integration events of ~100-300.

**Isolation of single spirulina filaments**
From an actively growing spirulina culture, 200-500 filaments were spread on a SOT agar plate. The cells were allowed to settle on the plate for 1-2 h and examined under a dissection microscope. Well separated single filaments were picked with a 1 ml pipette tip and transferred in 3 ml SOT with appropriate antibiotics in around bottom tube. Typically, 10-20 single filaments were cultured in Innova for 15 days to propagate.

**Determination of transgene copy number**
To assess the copy number of an integrated transgene, 3 sets of primer/taqman probe pairs were designed to target 3 regions: an endogenous spirulina gene present at single locus (*cpcB*), a promoter region present at both an endogenous and transgenic locus (i.e. 2 chromosomal copies), and an exogenous region unique to the transgene. A synthetic g-block containing the 3 target loci plus flanking sequences was purchased from IDT as a calibrator. Real-time PCR was performed with the above primer/probes using genomic DNA from the transgenic spirulina strain and the g-block as templates. As controls, the parental spirulina strain and a second transgenic strain lacking template for the transgene-specific probes were tested. The relative copy number of the integrated transgene was calculated as the fold difference between the transgene and endogenous gene with $\triangle\triangle C_t$ method. The experiment was repeated 5 times with 3 separate preparations of genomic DNA. The expected abundance ratio for the endogenous gene, promoter, and exogenous gene was 1, 2, and 1 respectively.

**Axenic strain isolation**
To establish axenic spirulina strains, cells were washed with SOT media on 10 µm filters to exclude small, unicellular bacteria, and single filaments were isolated from cells captured on the filter. Cells were grown to a density of 0.5-1 OD750/ml in an Innova incubator with proper antibiotics. Cells were pelleted from 5 ml cultures by centrifugation for 10 minutes at 1600x rcf. To maintain sterility, the following steps were performed in a laminar-flow hood. The cell pellet was resuspended in 1 ml SOT and transferred to a 10 µm filter pre-wetted with SOT medium. Medium was removed by gravity filtration. Cells were washed with successive 1 ml aliquots of SOT media until at least 200 ml of total media had pass through the filter. The remaining cells were resuspended with 0.5 ml of SOT and transferred to a sterile Eppendorf tube. Filaments were counted under a microscope as above, and 200-500 filaments were spread on a SOT agar plate. Single filaments were isolated as above. After ~15 days, 10 µL of culture was spread on LB plates without antibiotics. Plates were incubated for 3-5 days in a 37°C incubator. Filament cultures free of contaminants on the LB agar plates were then seeded in 10 ml SOT with 2.5 g/L dextrose at a density of 0.1 OD750/ml. Cultures were grown in Multitron for 3 days. A 100 µL sample of the culture was plated on LB agar plates without antibiotics and incubated at 37°C for 5 days. Cultures with no contaminants observed on either set of LB plates were considered axenic.

**Culturing of non-spirulina microbes**
To culture non-spirulina microbes, a flask of spirulina culture was placed on bench for 3-5 h to allow the spirulina cells to settle down at the bottom of the flask. A 100 µL sample of supernatant was carefully pipetted and transferred to LB or mixed LB/SOT agar plates. Plates were incubated at 25-30°C on a light rack (60-70 µEi of light) for 5-7 days. Single colonies were streaked on the new plates for 5-10 rounds. Cells from single colonies were spread on fresh plates to propagate for further experiments.

**Identification of non-spirulina bacteria from spirulina cultures**
To culture non-spirulina microbes, a flask of spirulina culture was placed on bench for 3-5 h to allow the spirulina cells to settle down at the bottom of the flask. A 100 µL sample of spirulina-conditioned media was transferred to either LB agar, or mixed LB/SOT agar plates. Plates were incubated at 25-30°C on a light rack (60-70 µEi of light) for 5-7 days. Genomic DNA was extracted from bacterial samples following the genomic DNA extraction method described above. Highly conserved and degenerate 16S and 23S rRNA PCR primers and DNA was amplified as described[69,70] from samples derived from both LB and LB/SOT plates. PCR product libraries were subcloned and sequenced, revealing that sphingomonas was the dominant LB-derived isolate and microcella was the dominant LB/SOT isolate.

**Markerless strain engineering**
To create a platform for markerless integration, a parental strain containing a recombinant, non-native antibiotic marker was first generated. An integrating plasmid bearing homology arms for the D01030 (*kmR*) locus flanking an *aadA* gene for streptomycin resistance was transformed into wild-type spirulina. The integrating vector was designed to precisely replace the ORF of D01030 with the sequence for *aadA*. This vector was transformed into both UTEX (SP3) and NIES (SP7) spirulina strains, generating strains SP205 and SP207 respectively. After transformation, strains were propagated for 2 months and

confirmed to be fully segregated by genotyping. The strains were also challenged with kanamycin to demonstrate the loss of native kanamycin resistance.

**Verification of markerless spirulina strains**

Clonal isolates of fully segregated strains were verified as follows: 1) qPCR to demonstrate a single transgene per genome (see above); 2) sequencing of chromosomal DNA to verify the absence of mutations in the homology arms and inserted gene(s) (see above); 3) PCR to demonstrate loss of parental integration locus allele and complete segregation to homozygosity of the transgene (see above); 4) chromosome DNA sequence of the 16S rDNA locus to verify strain identity as *Arthrospira platensis* (see above); 5) sequencing of alternative insertion sites in chromosomal DNA to verify lack of strain contamination with other engineered spirulina strains (see above); 6) PCR to demonstrate absence of the integrating DNA vector backbone, which should be eliminated during integration by homologous recombination (see below); and 7) verification of streptomycin sensitivity and kanamycin resistance by antibiotic challenge.

The vector backbone sequences outside of the homology arms should not integrate into the genome and thus be absent from genomic spirulina DNA. To exclude the possibility of non-specific integration of the vector backbone DNA, PCR was performed with primer pairs targeting the ampicillin resistance gene and *E. coli* origin of replication. At no point have these fragments been observed in spirulina, suggesting that there is no integration of the vector outside of the homology arms.

**Construction of markerless transgenes for spirulina integration**

To ease cloning of transgenes into spirulina, a standardized vector was built for markerless integration. This “destination” vector included integrating homology arms for the *kmR* locus flanking an ORF for the native *kmR* gene and a terminator. The antibiotic marker was followed by a recombinant promoter/terminator pair for transgene expression. The promoter/terminator pair consisted of a constitutively active, native *A. platensis* promoter (600 bp upstream of the *cpcB* gene; named Pcpc600) and the terminator of the *E. coli* ribosomal RNA gene *rrnB* (named TrrnB). A pair of BsaI restriction endonuclease sites between the promoter terminator pair was used for Golden Gate cloning of protein coding sequences for transgenic expression. Protein coding sequences with compatible BsaI sites were purchased from IDT and cloned into the “destination” vector using a Golden Gate Assembly Kit (NEB). Plasmid DNA was purified from *E. coli* by QIAprep Spin Miniprep Kit (Qiagen) and transformed into the spirulina strain SP205. The product of integration of this construct is genetically identical to the wild-type *kmR* locus, excepting the transgene (i.e. no non-native antibiotic markers are present).

**Purification of recombinant protein from spirulina**

Recombinant aa682 was purified from spirulina by immobilized metal affinity chromatography (IMAC). Briefly, a 10 ml pellet of spirulina cells from strain SP1182 was collected from 2 liters of culture by centrifugation. Pellet was resuspended to a total volume of 35 ml with lysis buffer (50 mM sodium phosphate buffer pH 8.0, 500 mM NaCl, 20 mM imidazole) supplemented with Pierce Protease Inhibitor Minitablets (Thermo Scientific) and 1 mM phenylmethylsulfonyl fluoride (PMSF). The resuspension was passed through a french pressure cell twice to lyse the cells. Samples were kept on ice throughout. The insoluble fraction was pelleted by centrifugation at 5,000 x rcf for 30 min. The partially clarified lysate was mixed with 2 ml of washed HisPur Ni-NTA Resin (Thermo Scientific) and incubated at 4°C with gently rocking for 2 h. The resin was gently pelleted by centrifugation at 500 x rcf for 1 min., supernatant discarded, and resin resuspended in fresh lysis buffer. This process was repeated until the supernatant was clear. The resin was collected in a small column by gravity filtration, washed with 20 ml of lysis buffer, and spirulina-expressed aa682 was eluted with lysis buffer containing 200 mM imidazole. Purified aa682 was further polished by separation on a Superdex 75 Prep Grade column on an ÄKTA Pure, yielding a single band by SDS-PAGE electrophoresis.

**Preparation of spirulina extracts for analysis of soluble protein**

Soluble extracts from spray dried spirulina samples were prepared by a flash-freeze protocol. Dried spirulina biomass was resuspended in PBS containing Pierce Protease Inhibitor minitablets and 1 mM PMSF at a biomass concentration of 10-40 mg/ml in 1.7 ml Eppendorf tubes. Samples were mixed by inversion to resuspend biomass powder and flash frozen in liquid nitrogen for 2 min. Resuspensions were transferred to a 37°C water bath for 2-3 min. Samples were well mixed by inversion once thawed. The flash-freeze procedure was repeated 2 additional times. Biomass samples were then centrifuged at 15,000 x rcf at 4C for 30 min, and the soluble fraction was transferred to a separate tube for downstream applications.

**Expression analysis of recombinant proteins in spirulina**

Recombinant protein expression in spirulina was measured by capillary electrophoresis immunoassay (CEIA) using a Jess system (ProteinSimple). The Jess system was run as recommended by the manufacturer. Briefly, dried biomass samples were diluted to a concentration of 0.2 mg/ml using water and a 5X master mix prepared from an EZ Standard Pack 1 (Bio-Techne). Purified protein controls used to generate standard curves were typically loaded at a range of concentrations from 0.5-20 µg/ml. A 12-230 kDa Jess/Wes Separation Module (ProteinSimple) was used and 3 µL of each sample was loaded for 9 s. A mouse anti-His-Tag antibody (GenScript) was diluted 1:100 and used as the primary detection antibody. An anti-mouse HRP-conjugated secondary antibody (ProteinSimple) was primarily used for chemiluminescent detection; fluorescently labelled anti-mouse antibodies (ProteinSimple) for IR or NIR fluorescence detection were used for some experiments. Data analysis was performed using the Protein Simple Compass software.

**Expression, purification, and biotinylation of *E. coli* expressed proteins**
Recombinant *C. jejuni* flagellin was expressed and purified from *E. coli*. A region of flaA (Sequence ID: WP_178888959.1) predicted to be soluble and exposed on the surface of flagella (amino acids 177-482) was cloned onto the C-terminus of MBP in a pET28b *E. coli* expression vector. The vector was transformed into BL-21(DE3) cells were grown overnight at 37°C on agar plates with kanamycin, and a single colony was used to inoculate a culture of LB media containing kanamycin. Cells were grown overnight with shaking at 225 rpm at 37°C, back-diluted to OD600 of 0.05, and grown at 37°C until the cells reached mid-log phase (OD600 = 0.4-0.6). Cells were induced with IPTG and incubated with shaking at 16°C overnight. The following day, the cells were pelleted by centrifugation at 3,500 for 20 min at 4°C, resuspended in 30 ml of lysis buffer containing protease inhibitors, and lysed with a Q700 Sonicator (Qsonica). The MBP-flaA fusion was purified from the clarified lysate using Amylose Resin (NEB) per the manufacturer's recommendations, and purified protein was aliquoted and stored at -80°C. Biotinylated MBP-flaA protein was prepared using an EZ-Link NGS-PEG4-Biotin kit (Thermo Scientific) following the manufacturer's guidelines.

VHHs expressed in *E. coli* used similar expression vectors and bacterial cells lines. Culturing, induction, and lysis of *E. coli* expressing VHHs followed the same protocol as with flaA expression. Purification of the VHHs from lysates was performed by IMAC, following the purification protocol described for aa682.

The RBD antigen used with VHH-72 was a kindly provided by the Roland Strong (Fred Hutchinson Cancer Institute).

**Binding assays for spirulina-expressed VHHs**
The EC50 binding activity of VHHs as purified protein and in spirulina lysate was measured by ELISA. High-binding 96-well plates (Greiner Bio-one or NUNC MaxiSorp) were coated with antigen by adding 100 µL of 1-5 µg/ml recombinant flaA antigen in carbonate-bicarbonate buffer (Sigma) to each well and incubating overnight at 4°C. Plates were washed 3 times with 300 µL PBS supplemented with 0.05% Tween-20 (PBS-T). Washed plates were blocked with 250 µL PBS-TM supplemented with 5% non-fat dry milk (PBS-TM) for 2 hours at room temperature. Blocking solution was discarded and 100 µL of sample containing VHH was added to each well. VHH samples were prepared by diluting purified protein or spirulina extracts with PBS-TM, and samples in a dilution series were serially diluted with PBS-TM. Samples were incubated at room temperature for 1 h to allow VHH binding to antigen. After incubation, plates were washed with 300 µL PBS-T 3 times. Wash was discarded, 100 µL of primary antibody diluted with PBS-TM was added to each well, and plates were incubated at room temperature for 1 h. A 1:10,000 dilution of either a mouse anti-His-Tag antibody (GenScript) or rabbit anti-camelid VHH antibody cocktail (GenScript) was used as the primary antibody. After incubation, plates were washed 3 times with 300 µL PBS-T, and 100 µL of a secondary antibody was added to each well. An HRP-conjugated goat anti-mouse antibody or HRP-conjugated donkey anti-rabbit antibody was used as the secondary antibody. Plates were incubated at room temperature for 30-45 min at room temperature. Plates were washed twice with PBS-T and once with PBS. Plates were developed using either a SeraCare KPL TMB Microwell Peroxidase Substrate System (Sera Care Life Sciences) or 1-Step Ultra TMB-ELISA Substrate Solution (Thermo Scientific) following the manufacturer's recommendation. Peroxidase activity was quenched after 5-10 min with 50 µL of 1 M HCl or 2 M sulfuric acid. Absorbance at 450 nm was measured on an M2 plate reader (Molecular Devices). Data analysis was performed using Prism (GraphPad Software).

**Kinetics binding analysis of VHHs**
Kinetics binding measurements for were performed by biolayer interferometry (BLI) using an Octet Red96e (Forte Bio). Biotinylated MBP-flaA was loaded onto streptavidin biosensors with a loading concentration of 100 nM and loading time of 4 min. After loading, probes were allowed to reach a baseline equilibrium in kinetics buffer (PBS with 1% bovine serum albumin and 0.05% Tween-20) for 2 min. Association and dissociation were monitored for 20 s and 140 s respectively. Purified aa682 diluted with kinetics buffer was assayed at concentrations from 1 µM to 10 nM. The 10 nM sample was excluded from analysis for weak signal. Two biosensors were used as references: a 0 nM aa682 control, as well as a no ligand control. Kinetics binding values were determined using Octet Data Analysis HT software (ForteBio). Curve fits were performed using a global fit across all concentrations of aa682 and assuming a 1:1 binding model.

**Epitope mapping of VHH/antigen interaction**
The epitope mapping of the interaction between FlagV6 and flagellin was performed using phage displayed peptide fragments derived from a ~300 amino acid soluble fragment of *C. jejuni* flaA. A sliding window of 30 amino acid fragments, with a 2 amino acid interval along the length of flaA, was designed as oligos for cloning into the phagemid pADL-23c

(Antibody Design Labs). The peptide library was cloned into the BglI site of the phagemid by Gibson Assembly and transformed into DH5a *E. coli*, yielding >6 x $10^4$ transformants. The phagemid library was cleaned up by QiaPrep Spin Minikit columns and transformed into electrocompetent TG1 cells (Lucigen). Phage production was induced with the pIII deficient helper phage CM13d3 (Antibody Design Labs) to ensure polyvalent display of the peptide epitopes. Phage from an overnight culture in 2xYT media was precipitated and washed following the manufacturer's protocol. Wells of an ELISA plate were coated overnight with 100 µL of 1 µg/ml FlagV6 VHH in carbonate-bicarbonate buffer, washed with PBS-T, and blocked with PBS-TM. The phage library was diluted with PBS-TM to a concentration of $10^{12}$ phage/ml and incubated at room temperature for 30 min. The phage were then panned for VHH binders by adding 100 µL of blocked phage to wells of the ELISA plate and incubating on a vibrating platform for 2 h at room temperature. Unbound phage were washed from wells with 6, 300 µL washed with PBS-T. Bound phage were eluted with low pH by adding 100 µL of 100 mM glycine, pH 2.0, incubating for 5 min with shaking. The elution buffer was neutralized with 40 µL of 2 M Tris, pH 7.5 and used to reinfect phage competent TG1 cells (Antibody Design Labs). The library amplification and panning were performed for 2 additional rounds. After the third round of panning, all phagemid-containing colonies were observed to contain the same peptide fragment by Sanger sequencing. Two independent replicates of the experiment yielded overlapping fragments that mapped to the D3 domain of flaA.

**Flow cytometry of VHH binding to *C. jejuni***

The binding of spirulina expressed VHHs to a pure culture of *C. jejuni* was measured by flow cytometry. An aliquot of lysate prepared from spray dried spirulina biomass was incubated with an equivalent volume of $10^7$ cfu/ml of *C. jejuni* 81-176 for 1 h at 4°C. After washing with PBS containing 2% FBSs, bacteria were incubated for 30 min with the anti-His-Tag antibody [iFluor647] (GenScript). Samples were washed with PBS containing 2% FBS, resuspended in 2% paraformaldehyde and acquired on LSR Fortessa flow cytometer (BD Biosciences) using FCS and SSC parameters in logarithmic mode. Data were analyzed using the FlowJo software (TreeStar) or FACS Diva software (BD Biosciences).

**Motility inhibition assay**

The motility-inhibiting activity spirulina-expressed aa682 was measured by the motility of *C. jejuni* through soft agar. All *C. jejuni* culturing was performed in a tri-gas incubator at 40°C under microaerobic conditions (5% $O_2$, 10% $CO_2$) unless otherwise stated. Glycerol stocks of *C. jejuni* were first streaked on Campy Blood Agar Blaser plates (Thermo Scientific) and grown for 48 h. Bacteria were then used to inoculate 0.4% soft agar Mueller-Hinton plates by stab and incubated for 48 h. A slice of agar from the leading edge of the motility halos was used to inoculate 10 ml of Mueller-Hinton broth. Liquid cultures were incubated under standard conditions for 72 h. A spot of 20 µL of 5 mg/ml of purified aa682 in PBS was added to the center of soft agar Mueller-Hinton plates and allowed to fully adsorb into the agar. VHH spots were inoculated with 1 µL of OD600 = 0.03 of *C. jejuni* from the liquid culture. A samples and controls were set up in triplicate. Plates were incubated under standard conditions. The diameter of the motility halos was periodically measured and used to calculate area.

**Mid-scale production of spirulina biomass for pre-clinical trials**

To prepare biomass for pre-clinical mouse trials, the scale of spirulina culturing was increased and harvested biomass was spray dried. Spirulina cultures were initially propagated in shake flasks in media based on the standard cyanobacterial SOT media in Multitron conditions. Shake flask cultures were used to inoculate airlift reactors, with media modified by partial replacement of sodium bicarbonate with sodium carbonate, such that initial culture pH was 9.8. Cells were grown at light levels between 500 and 2500 µmol/m$^2$/sec, with temperature maintained at 35°C. As the culture utilizes $CO_2$ and grows the pH rises, and $CO_2$ is added to the airlift stream to maintain pH between 9.8-10. Cultures were inoculated at a concentration of 0.1-0.5 g biomass per liter by ash-free dry weight, and harvested by filtration at 2-4 g/L.

To prepare for spray drying, the harvested biomass was rinsed with a dilute 0.1% trehalose solution (to remove excess media salts), concentrated again by filtration, and then spray dried in a centrifugal nozzle spray dryer. Feed rate, airflow, and inlet air temperature were controlled to maintain an outlet air temperature of 68-72°C at the powder separation hydrocyclone. Once collected from the hydrocyclone, the powder was sealed and stored in airtight, opaque mylar bags to prevent exposure to moisture or light. The powder is stored at room temperature.

Prior to use in animal trials, spirulina biomass was analyzed to confirm strain identity. Dried biomass was genotyped to confirm the presence of the correct transgene and the absence of contaminating sequences (see above). CEIA and ELISA binding assays (see above) were also performed to confirm expression and binding activity of the spirulina-expressed VHH.

**Prophylactic treatment of *C. jejuni* infection in 2 mouse models**

Two independent mouse models were used to test the efficacy of spirulina-expressed VHHs in treating *C. jejuni* infection.

In a pilot experiment with the first model of *C. jejuni* infection, mice were fed a zinc-deficient diet (dZD)[38] prior to challenge. Animals were maintained per institutional protocols and fed a regular diet with *ad libitum* water for 3 days. Animals were then started on the study diet for 10 days, after which water was replaced with water containing an antibiotic cocktail for 3 days to condition gut flora for *C. jejuni* colonization. Water was replaced with untreated, antibiotic-free water for 1 day prior to *C. jejuni* challenge. On day 0, mice were given an inoculum of $10^6$ live *C. jejuni* cells (resuspended in PBS), strain 81-176, by oral gavage. Food and water were provided *ad libitum* throughout. Mice were given 5 doses of a spirulina resuspension before and after challenge. Spray dried spirulina biomass was resuspended in PBS at a concentration of 10% (w/v). A 200 µL resuspension was delivered by oral gavage on days -1, 0, 1, 2, and 3, relative to challenge. Day-of-challenge dosing was administered 60 min. prior to challenge. Food and water were withdrawn 30 min. prior to treatment, then provided *ad libitum.* To assess efficacy, mice were monitored for symptoms of diarrhea, changes in weight, and bacterial shedding in stool. Weight measurements were made daily for 7 days. Stool samples were collected on days 1, 3, and 7 post-challenge.

A second experiment using the first model of infection involved a change of study diet and a reduced spirulina dose. Animals were fed a Regional Basic Diet (RBD) for 10 days, followed by 3 days of antibiotic treatment. Untreated water was provided for 1 day prior to *C. jejuni* challenge. On day 0, mice were given an inoculum of $10^6$ live *C. jejuni* cells (resuspended in PBS), strain 81-176, by oral gavage. Food and water were provided *ad libitum* throughout. Mice were given 3 doses of spirulina before and after challenge. On days -1, 0, and 1 relative to challenge, mice were orally gavaged with 200 µL of spirulina resuspension or control. Day-of-challenge dosing was administered 60 min prior to challenge. Food and water were withdrawn 30 min prior to treatment, then provided *ad libitum.* Spirulina resuspension was prepared at a concentration of 2% (w/v) in PBS. To assess efficacy, mice were monitored for changes in weight, bacterial shedding in stool, and levels of biomarkers in cecum. Weight measurements were made daily for 7 days. Stool samples were collected on days 2, 4, 6, 8, and 10 post-challenge. On day 11, the levels of lipocalin-2 (LCN-2) and myeloperoxidase (MPO) were measured in stool and cecal contents by ELISA (DuoSet ELISA Mouse Lipocalin-2/NGAL, R&D Systems).

In the second model of *C. jejuni* infection, mice were orally treated with a range of spirulina concentrations to identify the minimally effective prophylactic dose of therapeutic. Mice were housed, 5 per cage, under standardized conditions (20 ± 2ºC, 55 ± 8% relative humidity, 12 h light/dark cycle). Food and water were available ad libitum, and mice were monitored daily. Mice were pre-treated orally with 10 mg of vancomycin in 200 µL PBS at 48, 24 and 12 h prior to spirulina administration. A single 400 µL dose of spray dried spirulina resuspended in PBS was administered by oral gavage to mice 1.5 h before infection with *C. jejuni* 81-176 ($10^8$ cfu/200µL PBS). To monitor the efficacy, mice were observed daily, and stools were collected at 24, 48 and 72 h post infection. To monitor the pathogen load, stools were resuspended and plated on Mueller Hinton agar plates containing 10 µg/ml of vancomycin and trimethoprim.

The cecal polymorphonuclear neutrophils (PMNs) were measured by flow cytometry 72 h post infection. Mice were sacrificed, the caecum was removed, opened longitudinally, delicately separated by caecal content and washed twice with ice cold PBS. The caecum was digested twice with RPMI and EDTA 5 mM for 30 min at 37°C. The filtrated fragments were then digested in RPMI 5% FBS (fetal bovine serum), 1 mg/ml collagenase type II, 40 µg/ml DNase-I for 40 min. The filtered suspension, containing the caecum lamina propria cells, was centrifuged for 5 min at 1,500 rpm and resuspended in RPMI complete medium. Single-cell suspensions from caecal lamina propria were stained with labelled antibodies diluted in PBS with 2% FBS for 20 min on ice. The following mouse antibodies (mAbs) were used: APC conjugated anti-CD11b (1:200), PE conjugated anti-GR1 (1:200). Samples were acquired on an LSR Fortessa (BD Biosciences) flow cytometer. Data were analysed using the FlowJo software (TreeStar, Ashland, OR, USA) or FACS Diva software (BD Biosciences, Franklin Lakes NJ, USA).

The inflammation status of mice was evaluated by measuring faecal lipocalin-2 (LCN-2) levels in fecal supernatants by ELISA (DuoSet ELISA Mouse Lipocalin-2/NGAL, R&D Systems). Briefly, feces collected at sacrifice were resuspended at 0.01 g per 100 µL PBS, centrifuged for 10 min at maximum speed, and diluted before performing the ELISA per the manufacturer's instructions.

**Large-scale, continuous culturing of spirulina**

Spirulina cultures were grown at large scale (250 L) in airlift reactors following protocols similar to the mid-scale reactors described above. Cultures were inoculated into the same media described above for mid-scale cultures at a concentration of 0.1-0.5 g biomass per liter by ash-free dry weight, grown under identical temperature and pH controls, and harvested by filtration over stainless steel screens at 2-4 g/L. A portion of the harvested culture was used to inoculate serial cultures, and the remaining harvested biomass was used for spray drying as above. The dried powder was sealed and stored at room temperature in airtight, opaque mylar bags to prevent exposure to moisture or light.

Post collection, quality control of powder lots include determination of concentration of the 6x-his tagged protein using CEIA conducted on a Jess system (ProteinSimple). Specific ligand binding activity is determined on an Octet Red96e biolayer interferometry instrument (Forte Bio) using recombinant, biotinylated *C. jejuni* flaA protein attached to streptavidin coated biosensors. In addition, microbial characterization by USP <61> and <62>, and elemental impurities determined by USP <233>.

#### Long-term stability of dried spirulina biomass

Batches of SP1182 spray dried biomass were stored at room temperature and collectively assessed for binding activity by ELISA. Duplicate biomass samples from each batch were resuspended in PBS and lysed by freeze-thaw extraction and clarified by centrifugation. The binding activity of aa682 present in the lysates was determined by ELISA with a recombinant flaA antigen as described above. Purified aa682 was used to generate a standard curve for binding activity by linear regression. The standard curve was used to calculate the concentration of aa682 in the SP1182 lysates. The percentage of expected VHH activity was determined by normalizing the aa682 concentration in each lysate to an assumed concentration of 3% aa682 per unit biomass.

#### *In vitro* gastric protease digests of dried spirulina biomass

Spray dried SP1182 biomass was exposed to simulated gastric fluid (SGF) to determine the stability of aa682 present in spray-dried spirulina. A sample of spray-dried SP1182 biomass was resuspended in PBS at 30 mg/ml. This resuspension was diluted 1:30 with pre-chilled SGF (50 mM Citrate-phosphate buffer pH 3.0, 94 mM NaCl, 13 mM KCl, pH 3.5 with 2000 U/ml Pepsin (MP Biomedicals)) and incubated in a 37°C water bath. Protease activity was neutralized by adding 50 mM NaOH and 1 mM phenylmethylsufonyl fluoride (PMSF). Samples were pelleted by centrifugation at 14,000 RPM for 5 min. Biomass pellets were solubilized using 1X NuPAGE LDS Sample buffer to a final biomass concentration of 1 mg/ml and heated at 90°C on heat block for 10 min. A similar process was used to assess the stability of purified aa682, absent the centrifugation step.

The stability and activity of biomass-encapsulated aa682 after exposure to low pH, simulated gastric buffer was assessed by CEIA and ELISA binding assay. Spray-dried SP1182 biomass was resuspended in either 50 mM bicarbonate buffer or citrate-phosphate buffer, pH 3.0 with 1 mM PMSF. Samples were incubated in a 37°C water bath for 60 min. After incubation, biomass resuspensions were pelleted at 10,000 RPM for 5 min. The supernatant was transferred into fresh tubes and stored at 4°C. Pellets were resuspended in 1 ml of 50 mM bicarbonate buffer to a final biomass concentration of 30 mg/ml and incubated in 37°C water bath for 30 min. Resuspensions were treated to 3 cycles of flash freezing in liquid nitrogen followed by thawing at 37°C to extract soluble protein. After the last thawing, samples were pelleted using a refrigerated tabletop centrifuge for 30 min at maximum speed to separate soluble protein from insoluble cellular debris. The supernatant was used to measure the expression level and binding activity of aa682 by CEIA and ELISA respectively.

#### *In vitro* protease digests with intestinal proteases

To measure intestinal protease resistance, SP1182 lysates were digested with trypsin and chymotrypsin and VHH binding activity was assessed by ELISA. Total soluble extract was prepared from a resuspension containing 40 mg of dried SP1182 biomass per ml of bis-tris buffer (20 mM bis-tris, pH 6.0) by the freeze-thaw protocol described above. Two volumes of soluble extract were mixed with 1 volume of protease in bis-tris buffer and 1 volume of PBS to yield a final digest concentration of 0.1 or 0.01 mg/ml of trypsin or chymotrypsin with a reaction pH of ~6.5. Digests were performed at 37°C for 1 h with shaking at 900 rpm on an Eppendorf Thermomixer. Protease activity was neutralized by adding an equivalent volume of 2 mM PMSF and 2X Pierce Protease Inhibitor Mini tablets (Thermo Scientific) in PBS. Binding activity of VHH to recombinant flaA was measured by ELISA as described above.
